# Rapid Bayesian learning in the mammalian olfactory system

**DOI:** 10.1101/706200

**Authors:** Naoki Hiratani, Peter E. Latham

## Abstract

Many experimental studies suggest that animals can rapidly learn to identify odors and predict the rewards associated with them. However, the underlying plasticity mechanism remains elusive. In particular, it is not clear how olfactory circuits achieve rapid, data efficient learning with local synaptic plasticity. Here, we formulate olfactory learning as a Bayesian optimization process, then map the learning rules into a computational model of the mammalian olfactory circuit. The model is capable of odor identification from a small number of observations, while reproducing cellular plasticity commonly observed during development. We extend the framework to reward-based learning, and show that the circuit is able to rapidly learn odor-reward association with a plausible neural architecture. These results deepen our theoretical understanding of unsupervised learning in the mammalian brain.

## 1 Introduction

It is crucial for animals to infer the identity of odors, in situations ranging from foraging to mating [Li and Liberles, 2015]. While some odors are hardwired [Ishii et al., 2017], most must be learned. Learning, however, is particularly difficult, especially in natural environments where odors are rarely presented in isolation, most odors are presented a small number of times, and odor identities are rarely supervised. Nevertheless, animals can learn to associate an odor with a reward in a few trials [Staubli et al., 1987, Linster et al., 2002]. Our goal here is to elucidate the local plasticity mechanisms that orchestrate this rapid learning.

To make sense of odors, the olfactory system has to do two things. One is to learn the weights that mediate the mapping from odors to olfactory receptor neurons (OSNs); the other is to infer which odors are present given OSN activity. If the weights were known, the latter problem could be solved using approximate Bayesian inference [Grabska-Barwińska et al., 2017]. And in a supervised setting – a setting in which the animal is told which odors are present – the weights could be learned efficiently using recently proposed Bayesian approaches [Aitchison et al., 2017, Hiratani and Fukai, 2018]. In realistic settings, however, the weights are not known and learning is unsupervised. Nevertheless, we can combine the above methods to simultaneously learn the weights and infer the odors.

The approach is straightforward, at least in principle: when inferring which odors are present, average over the uncertainty in the weights; then use the inferred odors to update the estimates of the weights, and, importantly, decrease the uncertainty. As the estimates of the weights become more accurate, inference also improves. However, while straightforward, exact calculation of this learning process is intractable, meaning the time it would take to accurately perform the update is exponential in the number of odors. Consequently, we have to use approximate methods. Here we use a variational one [Beal et al., 2003], but others are possible.

Although inference is approximate, our model still leads to faster learning of olfactory stimuli compared to previously proposed sparse-coding based approaches [Olshausen and Field, 1997, Tootoonian and Lengyel, 2014, Kepple et al., 2018]. It also provides some insight into olfactory circuitry: it reveals the advantage, relative to the rectified linear transfer function [Nair and Hinton, 2010], of sigmoidal-shaped f-I curves typical of biological neurons [Oswald and Reyes, 2008, Poirazi et al., 2003], and it reproduces the reduction in neuronal input gain [zhang, 2004, Carleton et al., 2003] and learning rate [Nissant et al., 2009] commonly observed during development. In addition, it predicts that the learning rate of granule cells should decrease as they become more selective, and thus exhibit lower lifetime sparseness [Willmore and Tolhurst, 2001, Wallace et al., 2017], something that is possible (if difficult) to test experimentally. And finally, we extended our model to an odor-reward association task, and found that learning of a concentration invariant representation at the piriform cortex helps rapid odor-reward association.

While our approach gives us a model that is reasonably consistent with mammalian olfactory circuitry, it is not perfect. In particular, the architecture predicted by our approximate Bayesian algorithm does not perfectly match the architecture of the olfactory system. However, a plausible olfactory circuit based on our model, but with the addition of recurrent inhibition among piriform neurons [Bolding and Franks, 2018], still learns to perform reward-based learning quickly. These results suggest that even at the circuit level, approximate Bayesian optimization may underlie rapid biological learning. But at the same time, our study reveals its limitation when applied to a complicated system.

## 2 Results

### 2.1 Problem setting

Let us denote odor concentrations by a vector ***c*** = (*c*_1_, …, *c*_*M*_), where *c*_*j*_ > 0 if odor *j* is present and *c*_*j*_ = 0 if it is not. Here, we define an “odor” as something like the odor of apple or coffee, not a single odorant molecule. In a typical environment, odors are very sparse, in the sense that a small number of them have a significant presence (i.e. *c*_*j*_ > 0 for a small number of *j* at any time; Fig. 1 left). In both vertebrate and invertebrate olfactory systems, odors are first detected by olfactory receptor neurons (OSNs), and then transmitted to glomeruli as spiking activity [Wilson and Mainen, 2006]. Neural activity accumulated at a glomerulus, denoted *x*_*i*_ for *i*-th glomerulus (and thus *i*-th OSN receptor type), is given approximately by

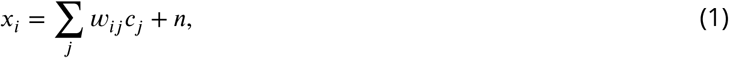

where *n* is the noise due to sensory variability and unreliable OSN spiking activity, and the weight, *w*_*ij*_, determines how strongly odor *j* activates glomerulus *i* (Fig. 1 right). OSN activity shows a roughly logarithmic dependence on odor concentration [O’Connell and Mozell, 1969, Hopfield, 1999]. Thus the amplitude, *c*_*j*_, of each zdor reflects log-concentration, not concentration. Below a threshold, here taken to be zero for convenience, odors are considered undetectable.

**Figure 1:**
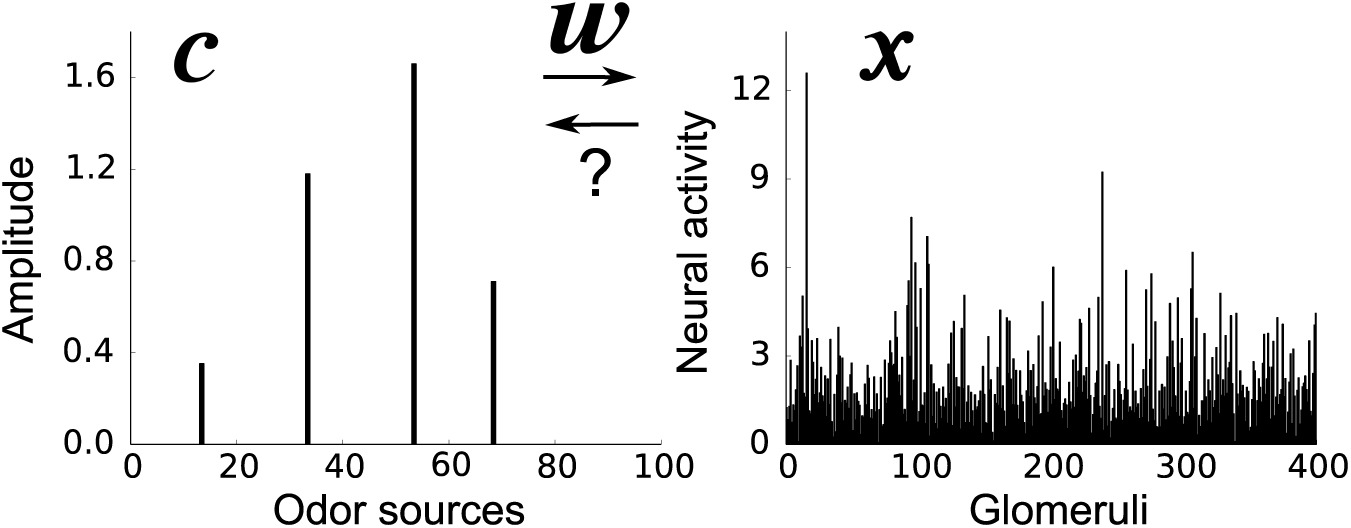
An example odor stimulus, ***c*** (left), and the response at the glomeruli, ***x*** (right). The weights, ***w*** (which are unknown to the animal) map odors, with concentration ***c***, to OSN activity accumulated at the glomeruli, ***x***. The goal of the animal is to infer the odor concentrations from the OSN activity.

### 2.2 Sparse coding

The goal of the brain is to infer which odors are present and what their concentrations are, based on OSN activity, ***x***. This is hard because the animal does not know the mixing weights, ***w***, but instead has to learn them, without supervision. One common approach to this type of unsupervised learning is the sparse coding model. Sparse coding was first proposed as a model of the early visual systems [Olshausen and Field, 1997], but it has been applied to olfaction as well [Tootoonian and Lengyel, 2014, Kepple et al., 2018]. The basic idea is that the odor, denoted ***ĉ***, and the weight matrix, denoted ***ŵ***, that best explains the input, ***x***, should be close to the real ***c*** and ***w***. This means ***ĉ*** and ***ŵ*** can be estimated by performing stochastic gradient descent on the likelihood of the inputs, ***x*** (see *Sparse coding model* in Methods). However, this is sub-optimal, primarily because uncertainty in ***ĉ*** and ***ŵ*** are ignored, even though they are important for data efficient learning [MacKay, 1992]. In addition, for tractability, the prior over the odors is taken to be a continuous function, making it difficult to capture the fact that at any given time most odors are absent. These constraints make the learning algorithm inefficient, and thus slow, as we show below (see in particular Fig. 4). For that reason, we turn to Bayesian inference.

### 2.3 Olfactory learning as Bayesian inference

The Bayesian approach is efficient because it allows uncertainty in both ***c*** and ***w*** to be taken into account, and it can naturally incorporate a prior that reflects the sparseness of the olfactory environment. The steps are straightforward: first write down, from Eq. (1), an expression for *p*(***c*|*x, w***), the distribution over odor concentrations given glomeruli activity, ***x***, and weights ***w***; then marginalize over the distribution of the weights given all the previous inputs, *p*(***w*|**past observations of ***x***) (see Methods, Eq. (10)). However, the exact marginalization is neither computationally tractable nor biologically plausible. We therefore employ a variational Bayesian approximation [Beal et al., 2003]. The basic idea is to replace the true joint probability distribution with a simpler one; here we use a fully factorized distribution, which makes marginalization especially easy.

The derivation of the algorithm for variational inference is described in detail in Methods. Here we simply give the results. The variational probability distribution of the concentration of odor *j* is updated iteratively as (see Methods, Eq. (14b))

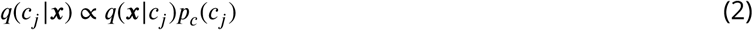

where *q*(***x*|***c*_*j*_) is the variational likelihood of the concentration of the *j*^th^ odor, *c*_*j*_, given ***x*** and *p*_*c*_ (*c*_*j*_) is the prior distribution over *c*_*j*_. We take the noise, *n*, that appears in Eq. (1) to be Gaussian, so *q*(***x*|***c*_*j*_) is Gaussian (Fig. 2A left). And to reflect sparsity, *p*_*c*_ (*c*_*j*_) is taken to be a point mass at zero combined with a continuous piece at positive concentration (Fig. 2A middle). Because, the prior strongly favors the absence of odors, the estimated mean concentration, ⟨*c*⟩ _*q*(*c*|***x***)_ (dashed black line in the right panel of Fig. 2A), is typically smaller than the mean over the likelihood function, ⟨*c*⟩_*q*(***x*|***c*)_ (dashed orange line in the right panel of Fig. 2A).

**Figure 2:**
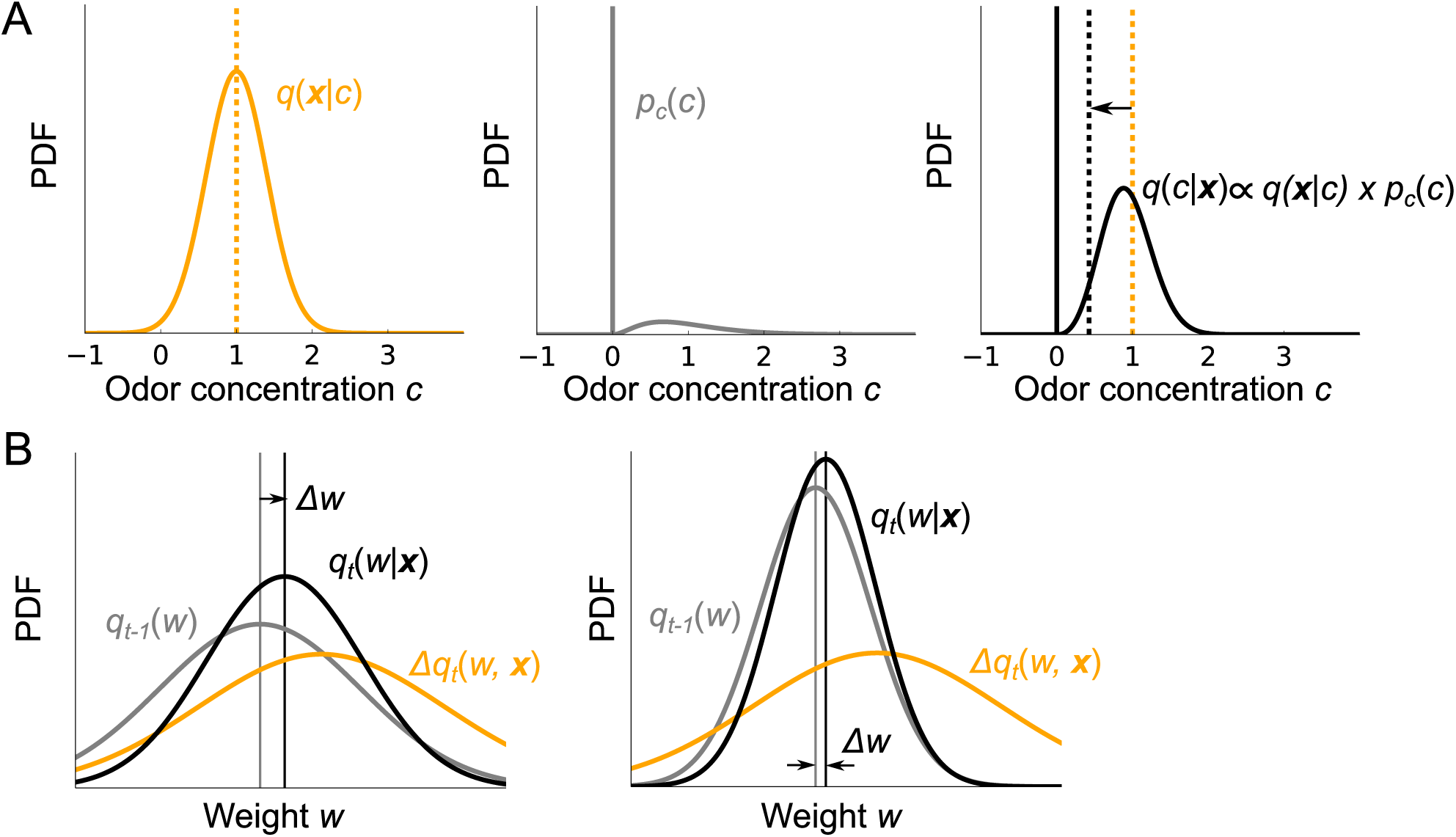
**A)** Inference of odor concentration. Combining the likelihood *q*(***x*** *c*) (left) and the prior *p*_*c*_ (*c*) (middle), the posterior distribution *q*(*c*|***x***) is obtained (right). The orange dashed line is the mean concentration associated with the likelihood, *q*(***x***|***c***); the black dashed line is the mean associated with the posterior, *q*(***c***|***x***). Because the prior strongly favors the absence of odors, the latter is shifted to lower concentration. **B)** Illustration of the weight update given the same sensory evidence b.*q*_*t*_(*w*, ***x***) when the previously estimated probabilistic distribution of the weight *q*_*t*-1_(*w*) is broad (left), or narrow (right). Note that the mean of *q*_*t*-1_(*w*) is the same in both panels.

Similarly, the update rule for the variational probability distribution of a weight is given by

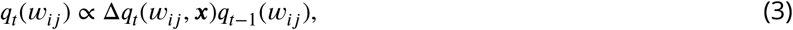

where Δ.*q*_*t*_(*w*_*ij*_, ***x***) is the evidence provided by the new information, carried in ***x***, at trial *t* (Fig. 2B) and *q*_*t*_(*w*_*ij*_) is the variational probability distribution of the weight, *w*_*ij*_, given observations up to trial *t* (a dependence we suppress to reduce clutter). Importantly, depending on the uncertainty in the weights, the same stimulus causes different amounts of plasticity, as illustrated in Fig. 2B. In particular, the higher the uncertainty in the estimated weight, *w*_*ij*_, at *t* - 1, the larger the change in the mean weight, Δ.*w* (left vs. right in Fig. 2B).

The update rules given in Eq. (2) and Eq. (3) can be mapped onto neural dynamics and synaptic plasticity that closely mirrors the mammalian olfactory bulb (Fig. 3A and B). The firing rate dynamics obeys the equations

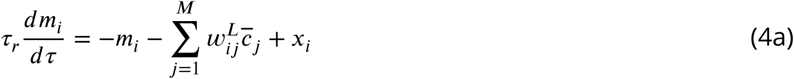

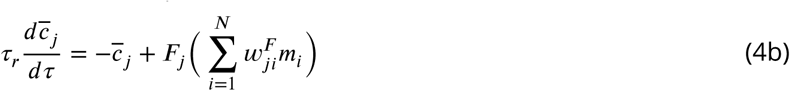

where *τ* denotes time within an odor presentation (not to be confused with *t*, which refers to trial), *m*_*i*_ is the firing rate of the *i*^th^ M/T (mitral/tufted) cell relative to baseline, and 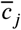 is the firing rate of the *j*^th^ granule cell. The *i*^th^ M/T cell is linearly modulated by excitatory inputs from glomerulus *i*, via *x*_*i*_, and also by inhibitory input from granule cells, the 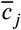. The granule cells are driven by excitatory inputs from M/T cells, mediated by a nonlinear transfer function *F*_*j*_. As we discuss below (see in particular Fig. 5), this nonlinearity plays a critical role in rapid learning. The activity of the *J*^th^ granule cell, 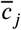, corresponds to the expected concentration of the odor represented by the cell.

**Figure 3:**
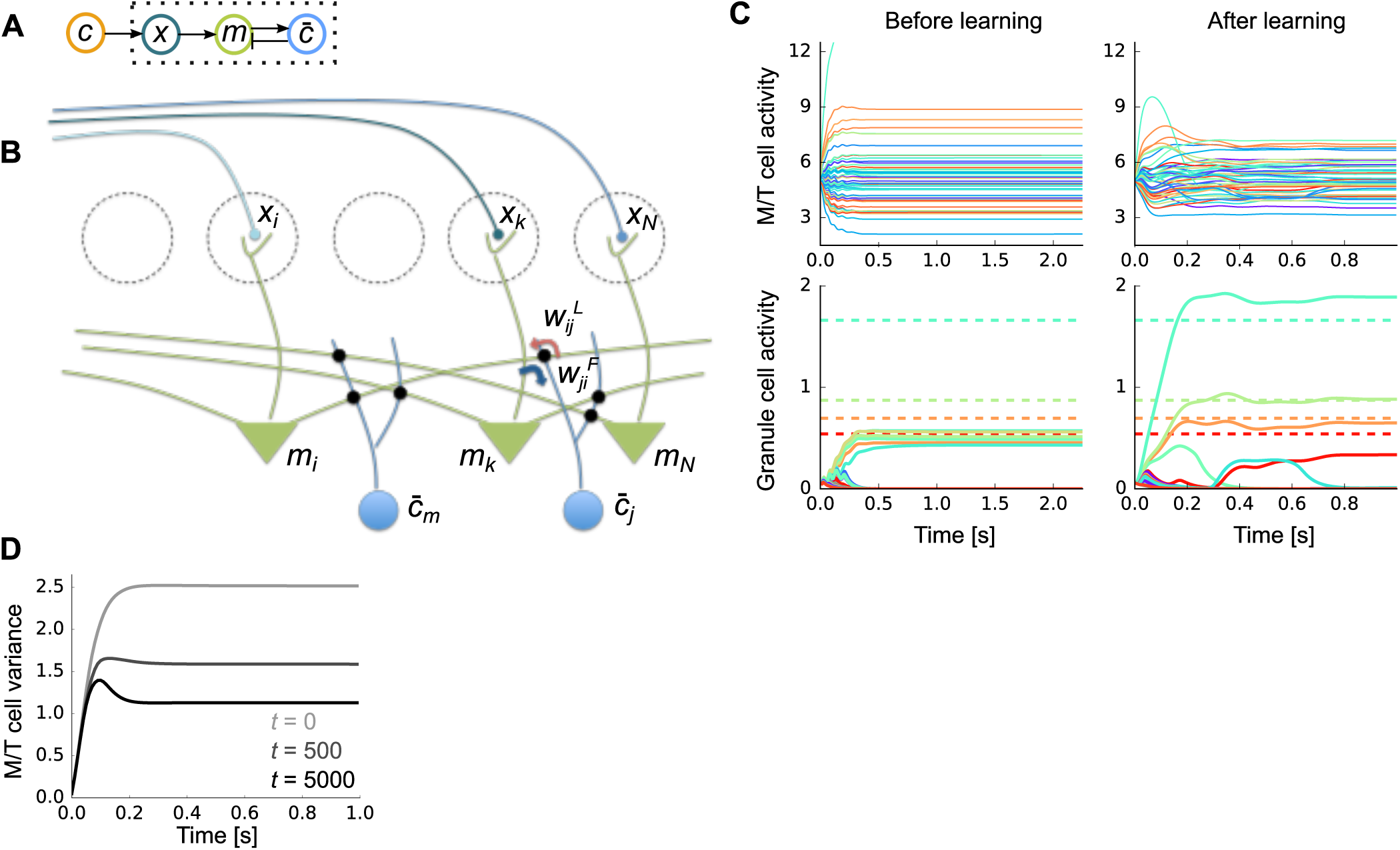
**A)** Schematic of the neural architecture. Dotted box represents the internal variables of the brain; the odor, ***c***, comes from the outside world. **B)** The neural implementation of our Bayesian learning model maps almost perfectly onto the circuitry of the olfactory bulb. Dotted circles are glomeruli, green triangle neurons are M/T cells, and blue neurons represent olfactory granule cells. **C)** An example of firing rate dynamics before (left) and after (right) learning (*M* = 50 odors, *N* = 400 glomeruli, and 4 odors were presented). Dotted horizontal lines in the bottom figure represent the true concentration of the presented odors. **D)** Change in the variance of M/T cells during learning (*t*: trial). The expectation was taken over both population and trials.

**Figure 4:**
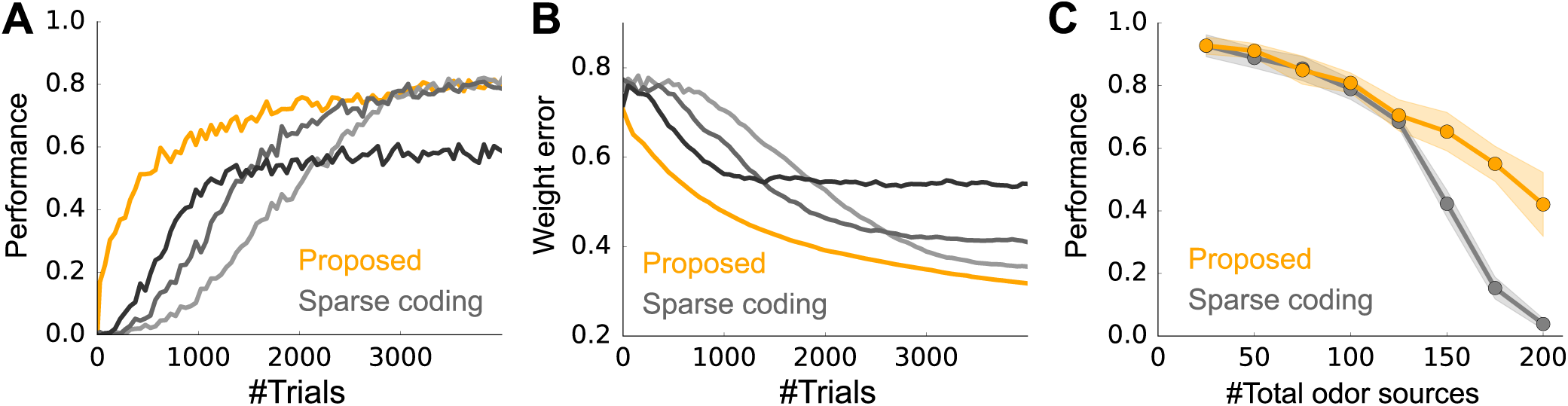
**A)** Learning curves for our model (orange) and sparse coding (grey-black). (*M* = 100 odors, *N* = 400 glomeruli, and on average, 3 odors were presented at each trial.) See *Performance evaluation* in Methods for details. The learning rates of the sparse coding model, *1*_*w*_, were 0.3, 0.5, and 1.0 from light gray to black. **B)** Same as **A**, but the performance was evaluated by the error in the weight. **C)** Performance (after learning from 4000 trials) of the proposed Bayesian model (orange) and the sparse coding model (gray) versus the number of odors. As in panels **A** and **B**, *N* = 400 glomeruli and 3 odors were presented on average. Here, *1*_*w*_ was fixed at 0.5. Error bars are the standard deviation over 10 simulation.

**Figure 5:**
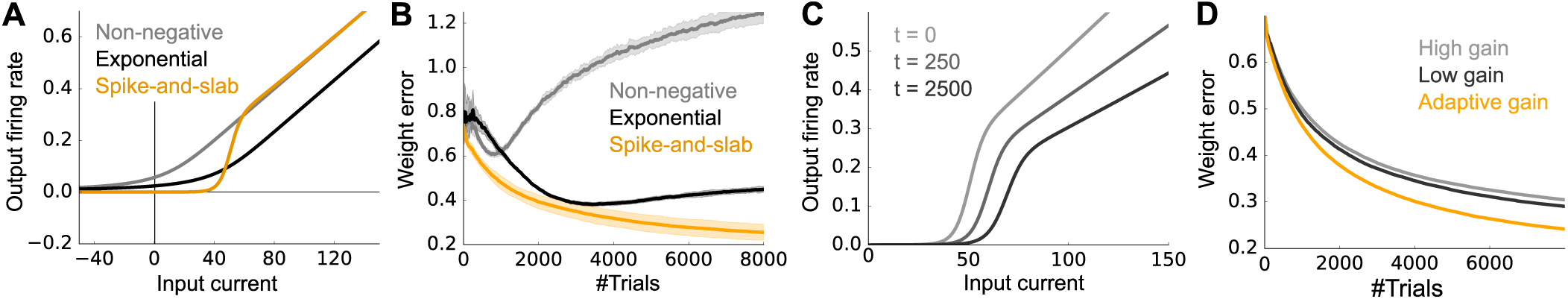
**A)** The shapes of the transfer functions of granule cells under different priors on the odor distribution. See Methods, *Models with various priors on odor concentration*, for the details. **B)** Weight errors under different priors. **C)** The average transfer function 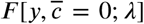 at the beginning (light gray; *λ* = 200), middle (gray; *λ* = 266), and the end (black; *λ* = 342) of the learning. The x-axis represents the input current *y*. **D)** The weight error under fixed input gains, compared to the control model with adaptive gain. In the gray (black) line, the transfer function was fixed to the top (bottom) curve in **C**. In all panels, *M* = 100 odors, *N* = 400 glomeruli, and 3 odors were presented on average.

The weights in Eq. (4), 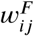 and 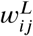, are the expected weights under our inference algorithm, represented by M/T-to-granule synapse and granule-to-M/T synapse, respectively (blue and red arrows in Fig. 3B). These synapses jointly form a dendro-dendritic connection between M/T and granule cells [Shepherd et al., 2007]. To keep track of the variational probabilistic distribution *q*_*t*_(*w*_*ij*_), both the mean and the variance of each weight need to be updated. The update of the mean is Hebbian, but with an adaptive learning rate,

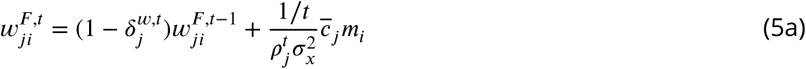

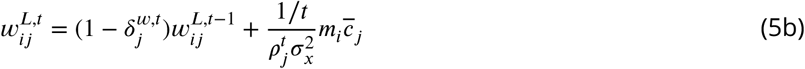

where *m*_*i*_ and 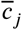 are evaluated at the end of the odor presentation. Here 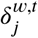 is the discount factor and 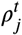 represents the precision (the inverse of the variance) of the synaptic weights 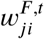 and 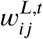(see *Synaptic plasticity* in Methods for details). This rule is Hebbian, as the update depends on the product of pre- and postsynaptic activity *m*_*i*_ and 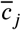. It’s also adaptive, as the update depends on the precision, 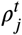: because of the 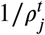 dependence, low precision (and thus high uncertainty) produce large weight changes while high precision (and thus low uncertainty) produce small weight changes. This is illustrated in Fig. 2B. The precision,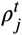, is also updated in an activity dependent manner; see Methods, Eq. (35).

Figure 3C describes typical neural dynamics before and after learning. Before learning, when a mix of four odors is presented, M/T activity quickly converges to constant values with a relative broad range (Fig. 3C, top-left), and granule cell activity is small and homogeneous (Fig. 3C, bottom-left). After learning, M/T cells exhibit transient activity, followed by convergence to a somewhat smaller range than before learning (Fig. 3C, top-right). Granule cells, on the other hand, show very selective responses, with activity levels roughly matching the concentration of the corresponding odors (Fig. 3C, bottom-right).

The activity profiles of cells in our model have many similarities with experimental observations. For instance, as observed in experiments [Gschwend et al., 2015], M/T cells show both positive and negative responses relative to baseline (Fig. 3C top, here the baseline is 5), and their responses are transient after some learning (Fig. 3C, top-right, and Fig. 3D). Moreover, the M/T cell responses become smaller as the animal learns the odors (Fig. 3D) as observed experimentally [Yamada et al., 2017]. In addition, after learning, granule cell activity is strongly modulated by odor concentration (Fig. 3C bottom-right; dotted horizontal lines represent the true concentrations of the corresponding odors), as observed experimentally [Tan et al., 2010]. The Bayesian approach is in principle optimal, but, given that we used an approximate model, how well did our circuit perform? We cannot compare to exact inference, because that is intractable. But we can compare to a close competitor: sparse coding. And indeed, our model does do better than the vanilla sparse coding model (Fig. 4). In particular, our model is able to learn much faster than the sparse coding model does (Fig. 4A). Moreover, to achieve high performance, the learning rate of the sparse coding model must be fine-tuned (gray lines in Fig. 4A). This advantage was replicated when we assessed the performance by the error in the weights (Fig. 4B). Despite faster learning, the asymptotic performance of the Bayesian model is similar to that of sparse coding when there are a relatively small number of odors in the environment, and much better when there are many odor sources, although the performance of both models deteriorates in that regime (Fig. 4C).

These results indicate that a variational approximation of Bayesian learning and inference enables data efficient learning, and does so using biologically plausible learning rules and neural dynamics. How does our model manage to perform fast and robust learning? And is there evidence that the brain uses this strategy? Below, we show that our proposed circuit performs well because it exploits the sparseness of the odors and utilizes the uncertainty in both the weights and odor concentration. We then discuss the relationship of our model to experimental observations.

### 2.4 The sparse prior leads to a nonlinear transfer function

An important feature of olfaction, like many real world inference problems, is that the distribution over odors has a mix of discrete and continuous components: an odor may or may not be present (the discrete part), and if it is present its concentration can take on a range of values (the continuous part). In our model, we formalize this with a so-called “spike and slab” prior (Fig. 2A middle): the “spike” is the delta function at zero; the “slab” is the continuous part. Sparseness is easy to enforce in this model: we just ensure that the cumulative probability of the slab, denoted *c*_*o*_, is close to zero. In most of our simulations, we used *c*_*o*_ = 3/*M* where *M* is the number of possible odors.

To see how the spike and slab prior affects the dynamics of the cells in our model, we note that the granule cells (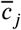 in Eq. (4)) represent the expected concentration of the odors, and so take the prior into account. Thus, after learning, most of them have near zero activity, with only a few of them active (Fig. 3C, bottom right panel). To achieve sparsity, the granule cells need a great deal of evidence to report non-negligible concentrations. That is reflected in the derived transfer functions of the granule cells (the function *F*_*j*_ in Eq. (4b); shown as the orange curve in Fig. 5A). The function exhibits near zero response (corresponding to near zero concentration) for small input, followed by an approximately linear response for large input.

If we derive update rules using a different prior, the transfer function changes. If we then perform inference and learning using the transfer function derived under a different prior, but drawing odors from the true prior, performance is, not surprisingly, sub-optimal (see Methods, *Models with various priors on odor concentration*). For example, if we constrain the odors to be non-negative (*p*_*c*_(*c*) ∝ Θ(*c*) where Θ is the Heaviside step function, an improper prior), the transfer functions are approximately rectified linear, a commonly used nonlinearity in artificial neural networks (gray line Fig. 5A; Nair and Hinton [2010]). However, this model failed to learn the input structure generated from the spike-and-slub prior, as the sparseness of the odor concentration is not taken into account (gray line in Fig. 5B). If we constrain the odors to be non-negative, but also ensure that they are not too large [Olshausen and Field, 1997], learning improves. In particular, if we use a prior of the form 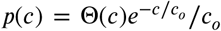, the transfer function is again approximately rectified linear, but it is shifted to the positive region compared to the curve for the non-negative prior (Fig. 5A; gray vs black lines). This enables the granule cells to have a weak response for a small input, resulting in successful learning, but convergence was slower than the model with the transfer function derived from the spike-and-slab prior (Fig. 5B). These results suggest that the classic input-output function – sigmoidal at small input and linear at large input – found both *in vitro* [Chance et al., 2002, Oswald and Reyes, 2008] and in biophysically realistic models of neurons [Poirazi et al., 2003], reflects the fact that the world is truly sparse – something not captured by classical sparse coding models. These gain functions thus offer a normative explanation for the biophysical responses of typical neurons to input. As a corollary, in early visual regions where the prior is arguably more continuous [Olshausen and Field, 1997], the rectified-linear transfer function might be more beneficial [Hennequin et al., 2018].

As the animal learns a better approximation to the true the weights, the olfactory system can extract more information from the OSN activity; this results in a change in the transfer function. In particular, the transfer function exhibits a decrease in gain with learning (mainly a shift to the right), as shown in Fig. 5C. Such a decrease in gain is a widely observed phenomenon among diverse neurons during development [zhang, 2004, Oswald and Reyes, 2008]. It is also consistent with the reduction of input resistance observed in adult-born granule cells during development [Carleton et al., 2003, Nissant et al., 2009], as a low resistance causes a low excitability. If the transfer functions were held fixed during learning, performance would deteriorate gradually (gray and black curves vs. orange line in Fig. 5D), though the benefit of the adaptive gain was rather small in our model setting.

The fact that the transfer function shifts to the right with learning seems counter-intuitive: the weights become more certain with learning, so it should take less input to the granule cells to produce activity, which suggests that the transfer functions should shift left, not right. However, an increase in certainty is not the only thing that changes with learning; the weights also become more diverse, capturing the diverse responses of glomeruli for each odor. The diversity increases the variance of the input to the granule cells, and so to ensure a sparse response with increasing diversity, the transfer functions need to shift to the right. In our model, increased diversity had a larger effect than increased certainty, resulting in a net rightward shift in the transfer functions, as shown in Fig. 5C (See Methods, *The variational odor distribution* for details).

### 2.5 Weight uncertainty leads to adaptive synaptic plasticity

A key aspect of our model is that it explicitly takes the uncertainty of the weights into account. This leads to an adaptive learning rate, as can be seen in Eq. (5). In particular, the learning rate is the product of two terms: 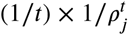. The first term, 1/*t*, is a global decay, and reflects an accumulation of information over time: at the beginning of learning, the olfactory stimuli contains a relatively large amount of information about the weights, and so the learning rate is large; at later times, the stimuli carry less information relative to what has already been learned, and so the learning rate is small. The second term, 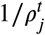, is the cell-specific contribution to the learning rate. In steady state, it is given approximately by 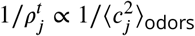 (the subscript “odors” indicates an average over odors; see Methods, Eq. (27a)).

It turns out that the second term can be related to the lifetime sparseness, denoted *S*_*J*_, which is defined to be 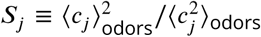 (see *Lifetime sparseness* in Methods and Willmore and Tolhurst [2001]). Assuming the mean firing rate, ⟨*c*_*j*_⟩_odors_, is approximately constant (as we see in our simulations), then 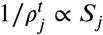. When the granule cells have broad, non-selective tuning, the lifetime sparseness is large, and the learning rate is high; when the cells are sparse and have highly selective tuning, the lifetime sparseness is low, and so is the learning rate. Thus, if the the mean granule cell responses are similar for all of the presented odors, the learning rate is large, encouraging neurons to modify their selectivity. If, on the other hand, the granule cell responses are sparse and selective, the learning rate is low, helping the neurons stabilize their acquired selectivity.

We examined the effects of the two factors – 1/*t* and 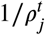 on learning. First, when the learning rate, 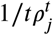, was kept constant throughout learning, learning was slower, even when the learning rate was finely tuned (gray lines vs orange line in Fig. 6A). This is consistent with the fine tuning required for the sparse coding model in Figs. 4A and B. When we fixed 1/*ρ*_*j*_ but included the global factor 1/*t*, performance was better than the model with fixed learning rate (light-green vs gray in Fig. 6A), yet still worse than the original fully adaptive model (light-green vs orange in Fig. 6A). This was more clear under a less sparse setting (*c*_*o*_ = 0.07, versus *c*_*o*_ = 0.03 in Fig. 6A), where the full model markedly outperformed the model with global decay (orange vs light-green in Fig. 6B). Furthermore, as predicted, we found that the learning rate of a cell, 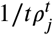, is positively correlated with the lifetime sparseness at each time point (i.e. at fixed *t*) as shown in Fig. 6D. This correlation becomes weaker as the prior becomes more sparse (compare Figs. 6C and D, for which *c*_*o*_ = 0.03 and 0.07, respectively). That is because a very sparse prior (low *c*_*o*_) helps the granule cells to be highly selective at an early stage, enabling the lifetime sparseness to quickly converge to a small value (vertical cluster on the left edge of Figs. 6C and D). These results indicate that the global and postsynaptic-neuron specific adaptation of the learning rate cooperatively help fast learning.

**Figure 6:**
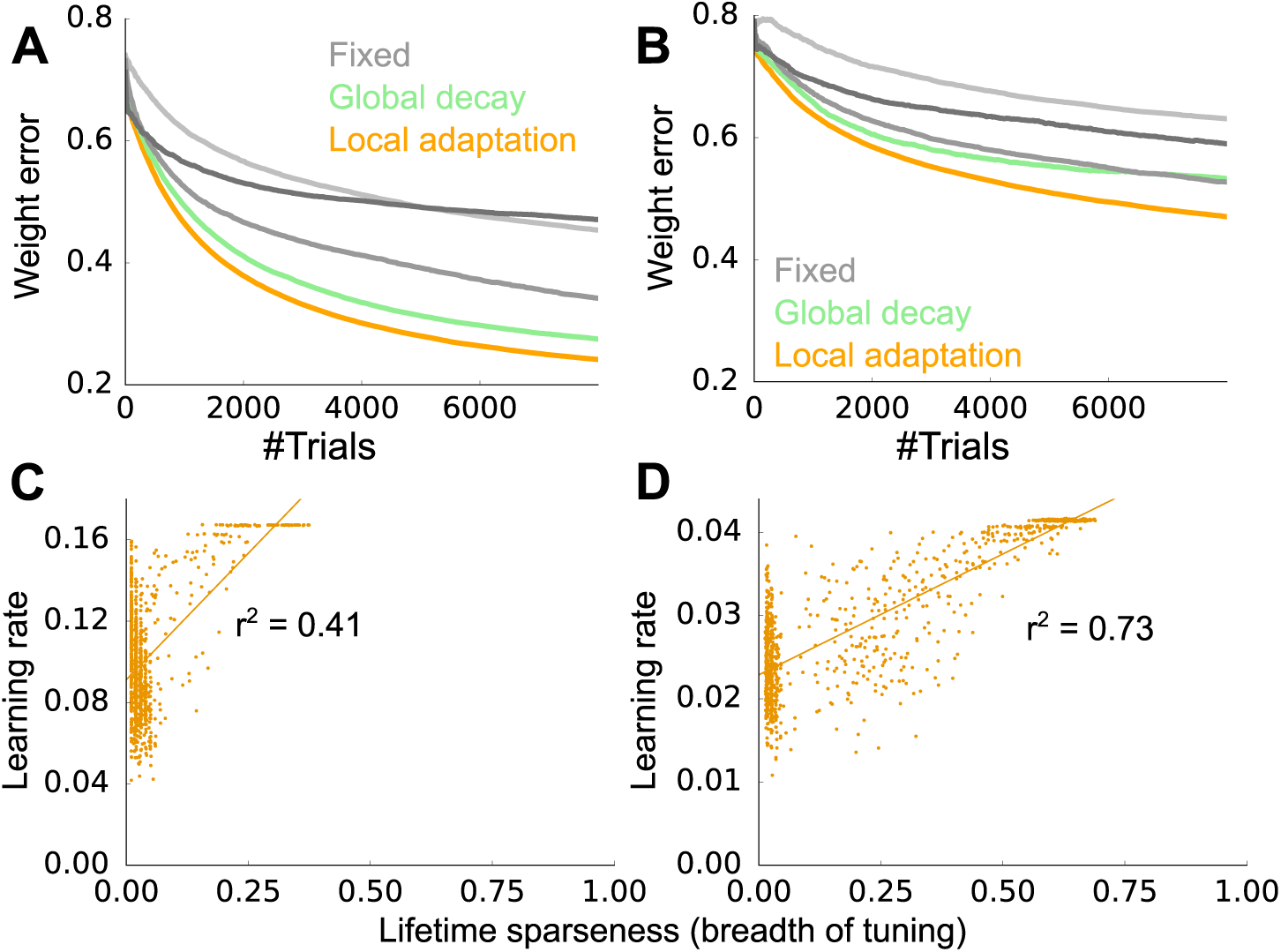
**A)** Weight error when 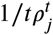 is fixed (gray lines), *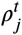* is fixed (light green), and fully adaptive (orange). For the gray lines we used learning rates of 0.01, 0.1, 1.0, correspond to light gray to dark gray. The sparsity, *c*_*o*_′ was 0.03. **B)** Same as panel **A**, but with a higher sparsity, *c*_*o*_ = 0.07. **C**,**D)** Correlations between the lifetime sparseness and the learning rate, after 300 stimuli were presented to the network, under more sparse (**C**: *c*_*o*_ = 0.03) and less sparse (**D**: *c*_*o*_ = 0.07) conditions. Lines are linear regressions, and each dot represents one granule cell. Vertical clusters appearing on the left edges of the panels correspond to neurons with very small lifetime sparseness. In all panels, *M* = 100 odors, *N* = 400 glomeruli, and 3 (**A**,**C**) or 7 (**B**,**D**) odors were presented on average.

### 2.6 Learning a concentration invariant representation and an odor-reward association

Our results so far indicate that olfactory learning is well characterized as an approximated Bayesian learning process. However, our circuit has a drawback: it estimates odor concentration, and not whether or not an odor is present. This is at odds with the demands of a realistic environment, where the existence of an odor is almost always far more important than its concentration. For instance, in a foraging situation, the perceived concentration depends on various factors such as the distance from the odor source, its size, and wind speed, and is almost independent of the associated reward. Thus, acquisition of a concentration-invariant representation is necessary for most olfactory-guided behavior.

From a probabilistic perspective, a concentration-invariant representation is essentially a representation of the probability of an odor being present, denoted 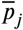. In our framework,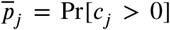; that probability is easily decoded from M/T cells using the circuit depicted in Fig. 7A(i) (see *Learning of concentration-invariant representation* in Methods). Here, 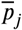 could be represented in layer 2 of piriform cortex neurons, as that is the the main downstream target of M/T cells, and odor representation in piriform cortex is approximately concentration-invariant [Bolding and Franks, 2018, Roland et al., 2017]. As the granule cells acquire odor representation, neurons in piriform cortex acquire odor probability representation (cyan and blue line in Fig. 7B(i)).

**Figure 7:**
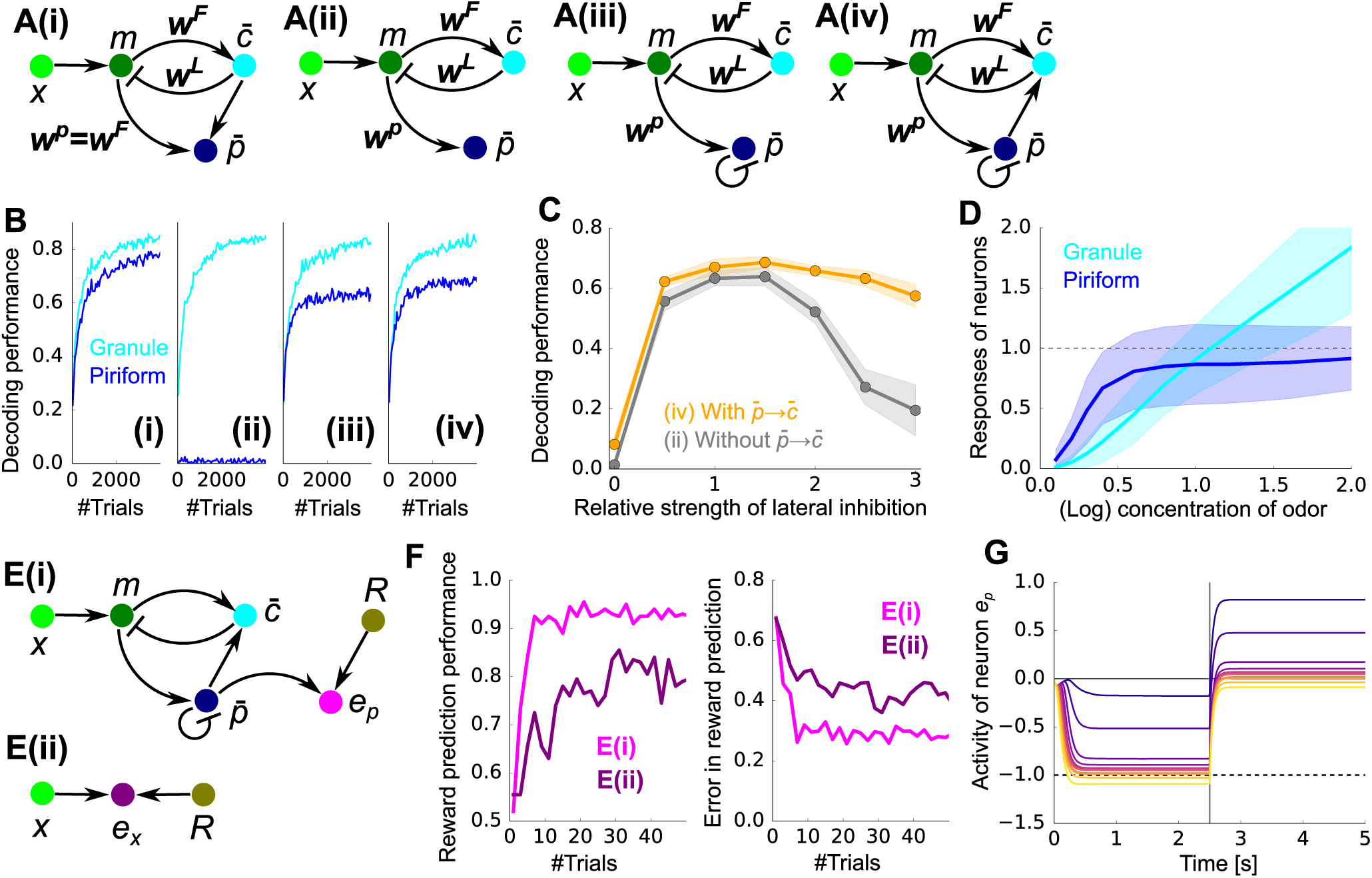
**A)** A set of increasingly realistic decoding models. **(i)** The decoding modal associated with our variational Bayesian inference algorithm. Note that the weights need to be copied from ***w***^*F*^ to ***w***^*p*^, something that is not biologically plausible. **(ii)** Similar circuit, but with the mapping from *m* to 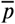 learned via a local rule. **(iii)** Same as **(ii)**, but with lateral inhibition. **(iv)** Same as **(iii)**, but with feedback to the granule cells. **B)** Learning performance for the models in **A. C)** Comparison of performance for model **A(iii)** (gray) and **A(iv)** (orange). **D)** Mean responses of the granule cells 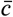 and the piriform neurons 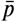 for their selective odors presented at various concentrations. The responses were measured by presenting each odor in isolation with different concentration, and then averaged over population. **E(i)** Schematic of the reward prediction circuit utilizing concentration-invariant representation in the piriform cells 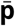. **E(ii)** Direct reward prediction from neural activity at glomeruli. **F)** Performance of odor-reward association measured by the classification performance (left) and the mean-squared error between the predicted reward and the actual reward (right). **G)** The mean response of neuron *e*_*p*_ given a odor associated with the reward. The vertical line at *τ* = 2.5*s* represents the reward presentation, and the dotted horizontal line is the sign-flipped reward value (-*R*). Different colors represents the different concentrations of the presented odor, from purple (*c* ≈ 0.1) to yellow (*c* ≈ 2.0). In all panels, *M* = 50 odors, *N* = 200 glomeruli, and 3 odors were presented on average, except for the go/no go task where one of two selected odors was presented randomly.

While the circuit shown in Fig. 7A(i) exhibits good performance, it is not consistent with what we know about the mammalian olfactory system, in two ways. First, the weights from the M/T cells to the granule cells have to be copied to the corresponding M/T to piriform cortex connections (i.e. ***w***^*p*^ = ***w***^*F*^), some-thing that is not biologically plausible. Second, a direct projection from granule cells to piriform cortex is needed, but such a connection does not exist. These inconsistencies can be circumvented by modifying the circuit heuristically (Fig. 7A(ii-iv)). Weight copying can be avoided by learning ***w***^*p*^ with local synaptic plasticity (Fig. 7A(ii)), although in the absence of the teaching signal from the granule cells, this naive extension does not work (blue line in Fig. 7B(ii)). However, introducing lateral inhibition among the piriform neurons, as observed experimentally [Bolding and Franks, 2018], allows the piriform neurons to acquire odor representation (Figs. 7A(iii) and 7B(iii)), although the decoding performance was worse than the Bayesian model (Fig. 7B(i) vs 7B(iii)). Here, lateral inhibition enables each piriform neuron to acquire different odor selectivity, but other mechanisms are possible. Finally, if connections from piriform cells to granule cells are added as well, the learning performance of granule cells became slightly better (Figs. 7A(iv) and 7B(iv)), and more robust to changes in the strength of lateral inhibition (Fig. 7C). As expected, the responses of piriform neurons were mostly concentration-invariant (blue line in Fig. 7D), whereas granule cells showed a clear concentration dependence (cyan line in Fig. 7D). Thus, the architecture of the mammalian olfactory circuit indeed supports robust learning of concentration-invariant representation, though somewhat different from the theoretical prediction.

Once the circuit acquires a concentration-invariant representation, constructing a circuit that performs odor-reward association is straightforward: all we have to do is take the circuit depicted in Fig. 7A(iv), and add a region that receives input from both piriform neurons and the reward system (*e*_*p*_ in Fig. 7E(i)). Olfactory tubercle could be the site for this odor-reward association [Wesson and Wilson, 2011], but it could be other regions, such as layer 3 of piriform cortex, as well. To test performance of this circuit, we implemented a go/no go task in which one odor is associated with a reward (*R* = 1.0), while another odor is associated with no reward (*R* = 0.0), regardless of concentrations. We simulated this task by randomly presenting rewarded or unrewarded stimulus with equal probability (see *Go/no go task* in Methods). We used the circuit pretrained with a large number of odors, but without reward. When the reward prediction was learned with the projection from piriform cells 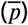 to olfactory tubercle cells (*e*_*p*_), classification performance reaches 90% after just 6 trials (Fig. 7F left). On the other hand, when the circuit learns the task directly from the glomeruli (Fig. 7E(ii)), though the circuit still learns to predict the reward as suggested previously [Mathis et al., 2016], learning was much slower and the performance was worse even after a large amount of training (purple vs pink lines in Fig. 7F). After a dozen odor-reward association from piriform neurons 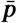, olfactory tubercle cell activity, *e*_*p*_, learned to represent the reward prediction given olfactory stimuli unless the concentration is very small (left half of Fig. 7G; in our model, -*e*_*p*_ is the reward prediction), and once the reward is presented at *τ* = 2.5*s*, the activity went back to near zero (right half of Fig. 7G; in our model, positive *e*_*p*_ represents an error, and so drives learning).

These results indicate that unsupervised learning of odor representation may underlie fast reward-based learning, and the proposed Bayesian learning mechanism improves reward association by enabling robust odor representation in a data efficient way.

## 3 Discussion

We formulated unsupervised olfactory learning in the mammalian olfactory system as a Bayesian optimization problem, then derived a set of local synaptic plasticity rules and neural dynamics that implemented Bayesian inference (Figs. 2 and 3). Our theory provides a normative explanation of the functional roles for the nonlinear transfer function and the developmental adaptation of the neuronal input gain (Fig. 5), both widely observed among sensory neurons. The model also predicts that the learning rate of dendro-dendritic connections should be approximately linear in the lifetime sparseness of the corresponding granule cells (Fig. 6). Finally, we extended the framework to learning of odor identity by piriform cortex, and showed that such learning supports rapid reward association (Fig. 7).

### 3.1 Relation to experimental observations

Our results suggest that adaptation of both input gain (Fig. 5) and learning rate (Fig. 6) are important for successful learning. The developmental reduction in input gain can be explained by a decrease in neural excitability, which is partially caused by the increased expression of *K*^+^ channels [Oswald and Reyes, 2008]. Correspondingly, it is known that changes in channel expression at the dendrite modulate the sensitivity of synaptic plasticity [Shah et al., 2010]. In particular, it has been reported that elimination of voltage-gated *K*^+^ channels enhances the induction of long term potentiation [Chen et al., 2006]. These results suggest that developmental up-regulation of *K*^+^ channel expression at the soma and the dendrite may underlie the adaptation of the input gain and learning rate.

The cellular plasticity rules we derived explain multiple developmental changes in adult born granule cells. Experimentally, relative to young cells, mature granule cells have sparser selectivity [Wallace et al., 2017], lower membrane resistance [Carleton et al., 2003, Nissant et al., 2009], and are less plastic [Nissant et al., 2009], as predicted by our model. In addition, our results provide insight into the functional role of adult neurogenesis. As shown previously [Hiratani and Fukai, 2018], if each synapse keeps track of its uncertainty, by removing the most uncertain synapses while adding synapses at a random position on the dendritic tree, a neuron can achieve sample-based Bayesian learning, making neurogenesis unnecessary. However, in our unsupervised learning framework, uncertainty is defined at neurons, not at synapses. As a result, from a Bayesian perspective, there is no good way to perform synaptogenesis. Thus, the brain should instead remove the most uncertain neurons, while at the same time randomly adding new ones.

### 3.2 Related theoretical work

The importance of the feedback circuit between M/T cells and granule cells has been noted previously [Koulakov and Rinberg, 2011, Grabska-Barwińska et al., 2017], but plasticity mechanisms that generate this circuit were not considered. Recently, several groups proposed learning algorithms for unsupervised olfactory learning using stochastic gradient descent [Beck et al., 2012, Tootoonian and Lengyel, 2014, Kepple et al., 2018], as in the case of our sparse coding model. However, as we have seen (Fig. 4), these algorithms are very unlikely to be fast. In addition to the sparse coding model, our problem setting is deeply related to Independent component analysis (ICA; Hyvärinen and Oja [2000]). Indeed, by using the sparseness as the measure of non-Gaussianity, unsupervised olfactory learning can be reformulated as an ICA problem [Tootoonian and Lengyel, 2014].

The spike-and-slab prior employed here is a widely used prior in machine learning [Mitchell and Beauchamp, 1988], and has been applied to the sparse coding model of the early visual system [Garrigues and Olshausen, 2008]. A contribution of this work is the establishment of a link between the spike-and-slab prior and nonlinear transfer function of a neuron. Triesch [2005] considered learning of hyper parameters that determines the shift and the steepness of transfer function while assuming a sigmoidal shape, and linked that with intrinsic homeostatic plasticity. However, the origin and the function of the sigmoidal transfer curves were not investigated.

Studies of adaptive learning rates, primally in the context of stochastic gradient descent, date back many decades [Amari, 1967, MacKay, 1992]; more recent studies have taken a Bayesian approach to adaptive learning in simplified single neurons models [Aitchison et al., 2017]. In this study, we considered an unsupervised learning problem, and showed that the learning rate of excitatory feedforward connections should depends only on the postsynaptic activity, independent of the presynaptic activity. Moreover, our theory predicted a non-trivial relationship between the learning rate and the lifetime sparseness of the postsynaptic neuron (Fig. 6C and D).

Acceleration of reward-based learning by unsupervised learning (Fig. 7F) has been studied in the context of both semi-supervised learning and model-based reinforcement learning. In particular, the latter approach has been applied to rapid learning by animals, but these were limited to abstract models, not circuit-based implementations [Doll et al., 2012]. In the invertebrate literature, Bazhenov et al. [2013] studied the combination of unsupervised and reward-based learning in a computational model of the insect brain, but plasticity was applied only to the output connections (corresponding in our model to *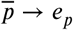* in Fig. 7E(i)). Interestingly, in the invertebrate brain, the connections corresponding to 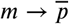 are mostly random and fixed [Caron et al., 2013], so the acceleration shown in Fig. 7F is potentially unique to vertebrates.

### 3.3 Limitation of the model

While our approach gave us a model that is reasonably consistent with mammalian olfactory circuitry, it is not perfect. In particular, the architecture predicted by our approximated Bayesian algorithm does not match perfectly the architecture of the olfactory bulb, piriform cortex, and olfactory tubercle (the latter involved in reward). We were able to make small modifications to our circuit so that it did match the biology, and still gave decent performance, but performance was not quite as good as the circuit predicted purely by Bayesian inference. On a broader scale, assuming our approach generalizes, it suggests that adult neurogenesis should exist wherever there is unsupervised learning, something that is not observed outside olfactory cortex [Ming and Song, 2011]. The discrepancy between the predicted and observed architecture highlights a limitation of this approach, especially when applied to complex systems. In particular, it is difficult to include biological constraints, both because we do not know exactly what they are, and because there is no straightforward way to marry those constraints to a normative Bayesian approach. However, that is an important avenue for future work.

## 4 Methods

### 4.1 Stimulus configuration

On each trial, the response of the *i*^*th*^ glomerulus is modeled as

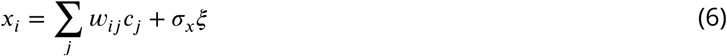

where *c*_*j*_ is the concentration of odor *j*, and ξ is a zero mean, unit variance Gaussian random variable. The Gaussian assumption is justified because, although olfactory sensory neurons fire with approximately Poisson statistics, 1000-10000 sensory neurons converge to a single glomerulus [Wilson and Mainen, 2006] where OSN activity is conveyed to M/T cells as stochastic currents. We take the weights, ***w***, to be log normal, followed by a normalization step,

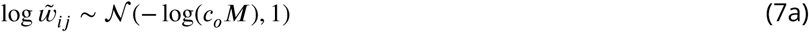

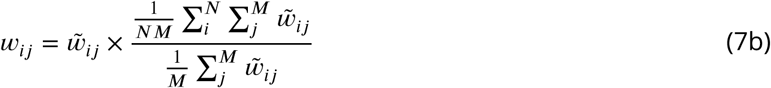

where, recall, *M* is the number of odors and *N* is the number of glomeruli. The factor multiplying 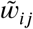 is 1 one average, so the normalization step doesn’t have a huge effect on the weights. However, it forces ∑_*j*_ *w*_*ij*_ to be strictly independent of *i*, which makes the learning process less noisy.

On each trial, odors *c*_*j*_ (*j* = 1, 2, …, *M*) are generated from the spike-and-slab prior given as

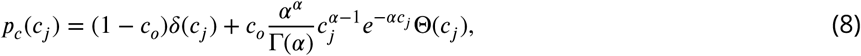

where Θ(*x*) is a Heaviside function. Throughout the manuscript, we used *α* = 3. Under this prior, each odor is independently presented with probability *c*_*o*_, and its amplitude follows a Gamma distribution with unit mean (Fig. 1A left). Note that the amplitude, *c*_*j*_, reflects log-concentration rather than the concentration [Hopfield, 1999]. To avoid the null stimulus, we resampled the odors if all of the *c*_*j*_ were 0 on any particular trial.

### 4.2 Bayesian model

The goal of the olfactory system is to infer the odor at time *t*, ***c***_***t***_, given all past presentations of odors, ***x***_**1:*t***_ **≡** {***x***_1_, ***x***_2_, …, ***x***_*t*_}. Because the weights are not known, they must be integrated out,

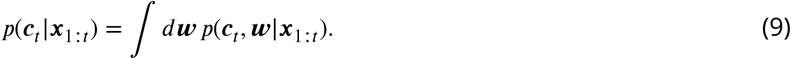

Using Bayes’ theorem, this can be written in a more intuitive form,

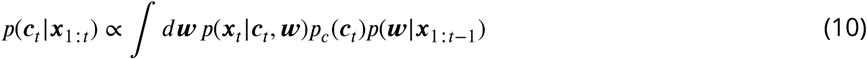

where, recall, *p*_*c*_(***c***_*t*_) is the prior over odors. To derive this expression, we used two facts: given ***c***_*t*_ and ***w, x***_*t*_ does not depend on past observations, and ***c***_*t*_ does not depend on past observations. The first term on the right hand side, *p*(***x***_*t*_|***c***_*t*_, ***w***) is the likelihood given the weights; but because we do not know the weights, we have to marginalize over them given past observations. The marginalization step, however, is intractable, as we have to introduce past odors and then integrate them out. While those integrals are tractable, the resulting distribution over ***w***, Eq. (10), cannot be performed analytically. And even if it could, the circuit would have to memorize all past stimuli, ***x***_1_, ***x***_2_,.., ***x***_*t*-1_. We thus have to perform approximate inference. For that we make a variational approximation.

#### 4.2.1 Variational approximation

The integral in Eq. (10) becomes easier if the distributions factorize. We thus make the variational approximation

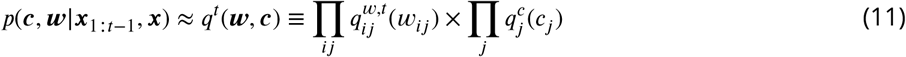

where, to avoid a proliferation of subscripts, we suppress the fact that ***c*** and 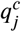 are to be evaluated at trial *t*; in line with this, to simplify subsequent equations we replace ***x***_*t*_ with ***x***; and, as is standard, we suppress the dependence of *q* on ***x***_1:*t*_.

The variational distributions, 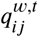 and 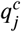, are found by minimizing the KL-divergence to the true distribution, with the KL-divergence given by

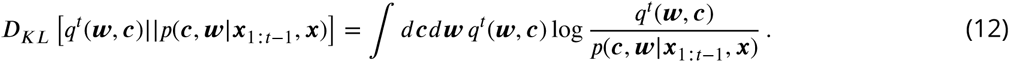

As is straightforward to show [Beal et al., 2003], minimizing this quantity leads to the update rules

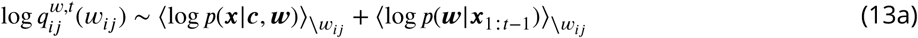

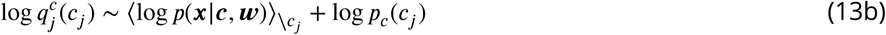

where ∼ indicates equality up to a constant, \*w*_*ij*_ indicates an average with respect to the variational distribution over all variables except *w*_*ij*_, and, similarly, \*c*_*j*_ indicates an average with respect to the variational distribution over all variables except *c*_*j*_. In the first equation, we approximate *p*(***w* |*x***_1:*t*−1_) with the variational distribution at the previous time step, 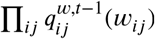, which makes the marginalization self-consistent. This approximation breaks down early in the learning process; nevertheless, in practice it works quite well. Using this approximation, we arrive at

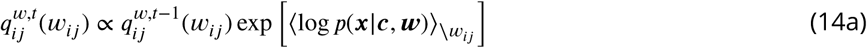

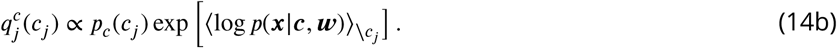

In the next two subsections we derive explicit update rules by computing the averages in these expressions.

#### 4.2.2 The variational odor distribution

To find the variational distribution over odors, we need to compute the average over log *p*(***x*| *c, w***) that appears on the right hand side of Eq. (14b). Using the fact that the ***x*** follows a Gaussian distribution, we have

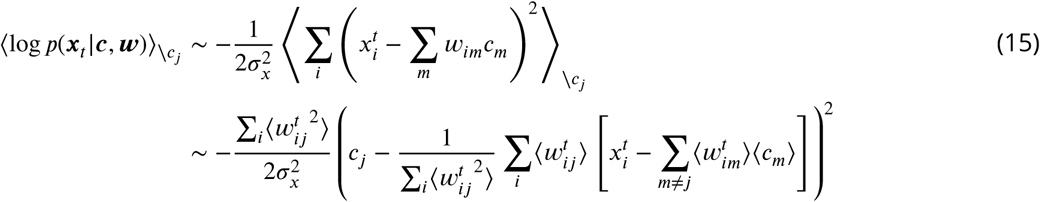

where the averages are with respect to the variational distribution. This is Gaussian, and it is straightforward to work out the mean and variance. Note that both depend on the first and second moments of the weights (which, as we will see below, determine the variational weight distribution) evaluated, importantly, at time *t*. However, synaptic plasticity is much slower than neural dynamics, so it is reasonable to update the weights on a slower timescale than concentration. Thus, when evaluating the mean and variance, we use the weight distribution on the previous time step. Using 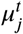 and 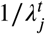 to denote the mean and variance, and making this approximation, we have

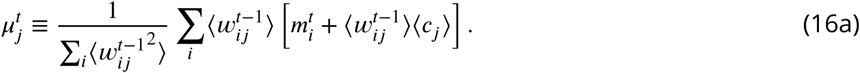

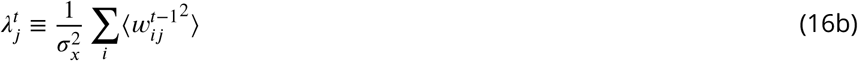

where we made the definition

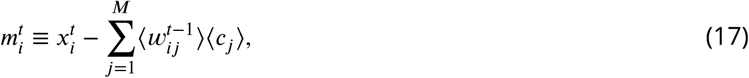

With these definitions,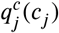 can be written in a very compact form,

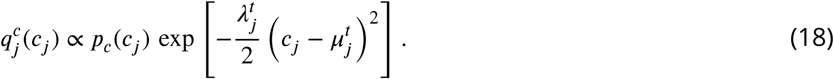

This is not a Gaussian under the spike-and-slab prior *p*_*c*_, which is not especially easy to deal with. However, as we will see below, to update the weights we do not need the full distribution; we just need the first and second moments. And for the reward-based learning, we need the probability that *c*_*j*_ is positive. These quantities are straightforward, if tedious, to compute.

For the mean, we have

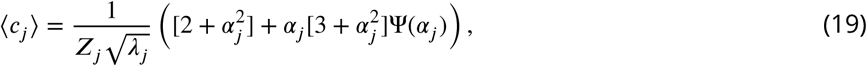

where the average is with respect to the distribution in Eq. (18), *Z*_*j*_ is the normalization constant

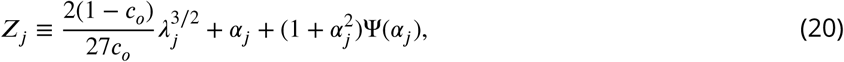

and *α*_*j*_ and Ψ(*α*) are defined by

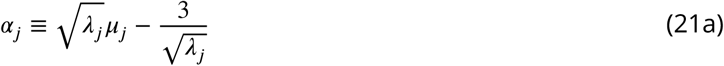

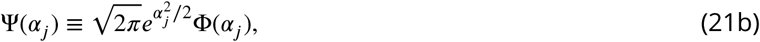

with Φ the cumulative normal function:

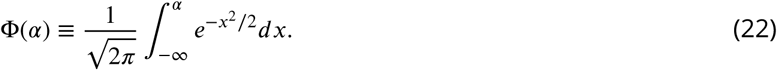

Similarly, the second moment is given by

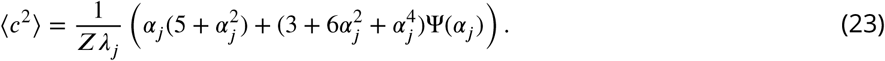

And finally, the probability that an odor is present is written

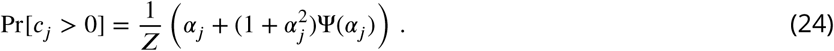

#### 4.2.3 The variational weight distribution

To find the variational distribution over weights, we need to compute the average on the right hand side of Eq. (14a). This is the same as Eq. (15), except that the average now excludes *w*_*ij*_ rather than *c*_*j*_,

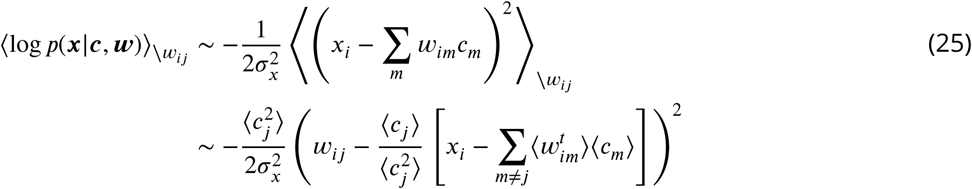

where the averages are, as above, with respect to the variational distributions. This is a quadratic function of *w*_*ij*_; thus, if we assume that 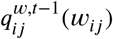 is Gaussian, then 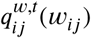 will also be Gaussian. Using 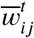 and 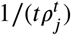 to denote the mean and variance at time *t*, respectively (the latter to anticipate the 1/*t* falloff of the variance expected under Bayesian filtering), Eq. (14a) becomes

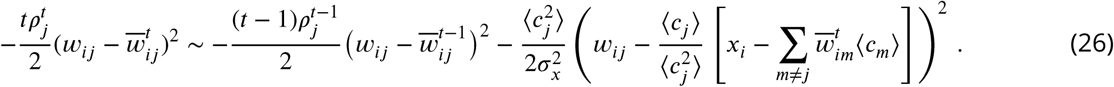

As in Eq. (15), 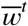 appears on the right hand side of Eq. (26). However, a very fast synaptic plasticity is required for solving this equation recursively for all the weights. We thus approximate the right hand side by using the previous timestep, *t* − 1, rather than the current one, *t*; an approximation that should be good when the weights change slowly. Doing that, we arrive at the update rules

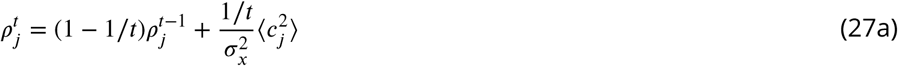

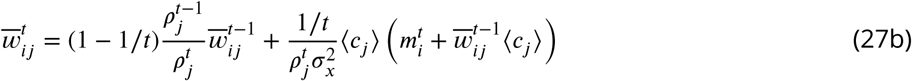

where we used Eq. (17) to simplify the second expression. Note that the update rule for 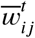 is local, as it depends only on variables indexed by *i* and *j*. The update rule for *ρ*_*j*_ is also local, and in fact depends only on variables indexed by *j*.

Finally, it is convenient to write the update rules for the mean and precision of the variational distribution over concentration, Eq. (16), in terms of 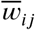 and *ρ*_*j*_,

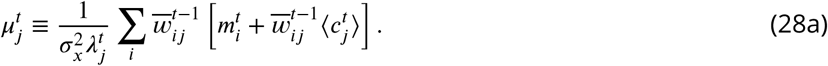

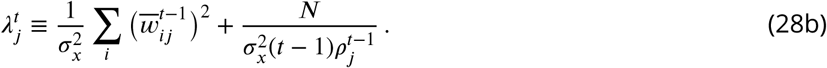

### 4.3 Network model

The analysis in the previous section revealed that under the variational approximation, the distribution of the odors and the weights are updated locally. Thus, we implement the update rules in a network model of the olfactory bulb. The update of the weight distribution,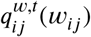, depends on ⟨*c*_*j*_⟩ and 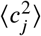, as shown in Eq. (27), while the update of the odor distribution,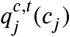, depends on 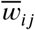 and ρ_*j*_, as shown in Eq. (28). Ideally, all these parameters should be updated simultaneously. However, as mentioned above, updates to synaptic weights are typically much slower than the neural dynamics, so here we consider a two step update. First, the relevant parameters of the variational odor distribution, ⟨*c*_*j*_⟩ and 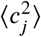, are updated using the mean and precision of the weight distribution, 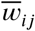 and *ρ*_*j*_, evaluated at *t* − 1. Then,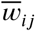 and *ρ*_*j*_ are updated using the first and second moments of the weights, ⟨*c*_*j*_⟩ and 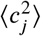, evaluated at time *t*.

#### 4.3.1 Neural dynamics

Our goal is to write down a set of dynamical equations for ⟨*c*_*j*_⟩ and 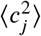 whose fixed points correspond to the values given in Eqs. (19) and (23), respectively. Examining these equations, we see that ⟨*c*_*j*_⟩ and 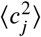 depend on *α*_*j*_ and *λ*_*j*_; after a small amount of algebra (involving the insertion of Eq. (28a) into (21a)), *α*_*j*_ may be written

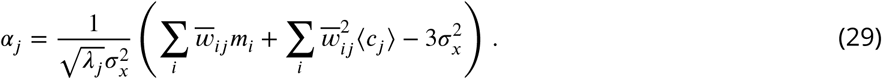

To avoid clutter, we have dropped the dependence on time, but the weights should be evaluated at time *t* – 1 and all other variables at time *t*.

Because neither *α*_*j*_ nor *λ*_*j*_ (the latter given in Eq. (28b)) depend on 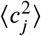, we can write down coupled equations for ⟨*c*_*j*_⟩ and *m*_*i*_; the solution of those equations gives us the values of *α*_*j*_ and *λ*_*j*_, which in turn gives us, via Eq. (23), 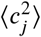. Using, for notational ease, 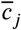 rather than ⟨*c*_*j*_⟩, the simplest such equations (derived from Eqs. (17) and (19)) are

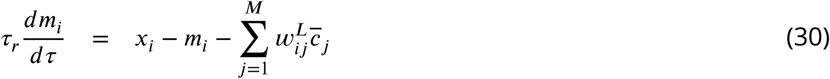

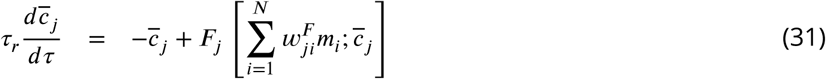

where *τ*_*r*_ is the time constant of the firing rate dynamics, and the nonlinear transfer function, *F*, is given by the right hand side of Eq. (19),

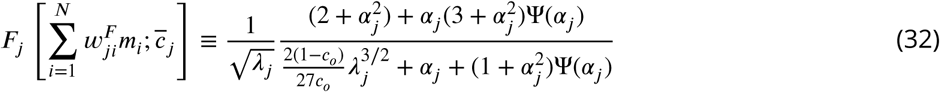

with *α*_*j*_ given in Eq. (29) and *λ*_*j*_ in Eq. (28b). Note that we have replaced the average weights, 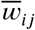, with two different weights, 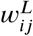 and 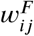. Ideally, we should have 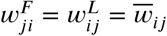, but, for biological plausibility, we allow that these reciprocal synapses to be learned independently. Note that when evaluating *α*_*j*_, Eq. (29), 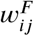 should be used. Although the expression for *F*_*j*_ seems complicated, the shape of the transfer function resembles the experimentally observed ones (see Fig. 5 below).

As shown in Fig. 3B, this dynamical system resembles the neural dynamics of the olfactory bulb, under the assumption that *m*_*i*_ and 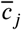 are the firing rates of M/T cells and the granule cells, respectively. With this assumption, 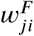 is the connection from M/T cell *i* to granule cell *j* and 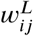 is the connection from granule cell *j* to M/T cell *i*.

Finally, the second moment of the concentration, is given, via Eq. (23), by

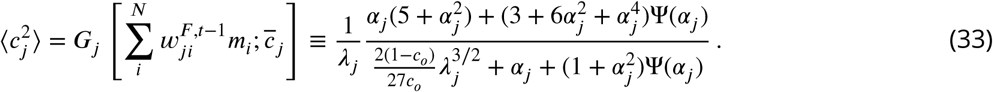

#### 4.3.2 Synaptic plasticity

After trial *t*, the average feed forward weights, 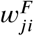, and the average lateral weights, 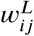, are updated as in Eq. (27b),

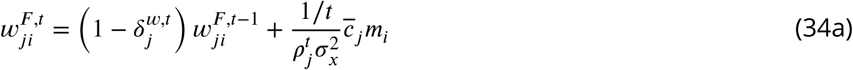

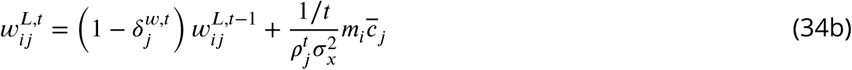

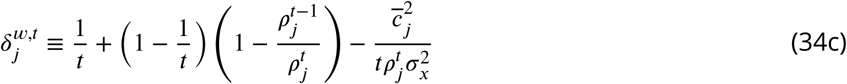

We used the firing rate *m*_*i*_ and 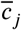 at the end of trial, after the neural dynamics has reached steady state. As the weight updates primarily depend on the product of *m*_*i*_ and 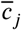, the learning rules are essentially Hebbian. Note that if the initial conditions are the same (i.e., if 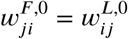), then 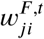 and 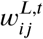 will remain the same for all time. This is reasonable given that connections between M/T cells and granule cells are dendro-dendritic.

The variance of the weights, 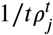, consists of two components. The first, 1/*t*, represents the global hyperbolic decay in the learning rate due to accumulation of information. In the simulations, we started *t* from *t* = *t*_min_ to suppress the influence of the initial samples. This is equivalent to using a trial-dependent discount factor 1/(*t*+*t*_min_) instead of 1/*t*, where *t* is the actual trial count. The second, 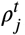 represents the neuron-specific contribution to the precision, and is given, via Eqs. (27) and (23), by

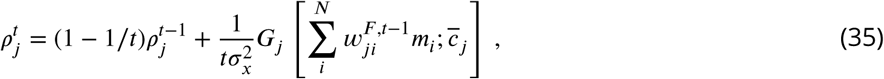

where, *G*_*J*_, the second moment of the concentration, is given in Eq. (33)

### 4.4 Models with various priors on odor concentration

In our model setting, the prior over concentration, *p*_*c*_(*c*), enters via Eq. (14b), and affects the transfer functions *F* and *G*, given in Eq. (32) and (33), respectively. Choosing different priors gives different transfer function. Below we consider two common ones.

#### 4.4.1 Non-negative

Because concentration is a trivially non-negative quantity, the random variable *c* should be non-negative as well. We can capture this with an improper prior, *p*_*c*_(*c*) ∝ Θ(*c*). This results in gain functions of the form

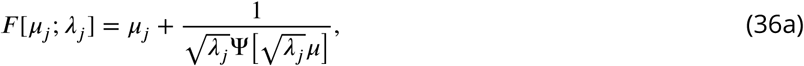

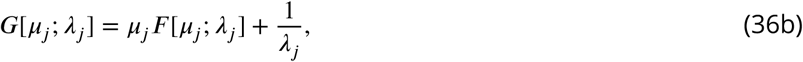

where *µ*_*j*_ and *λ*_*j*_ are given in Eqs. (28a) and (28b), respectively.

#### 4.4.2 Exponential

Under the non-negative prior introduced above, all positive concentrations are equally likely. However, that is not the case in a typical environment. Far more realistic is to assume that large concentrations are exponentially unlikely, yielding a prior of the form 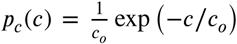. (The decay constant, *c*_*o*_, was chosen so that the mean is equal to *c*_*o*_, the same mean as in the true generative model.) For this prior, the functions *F* and *G* are

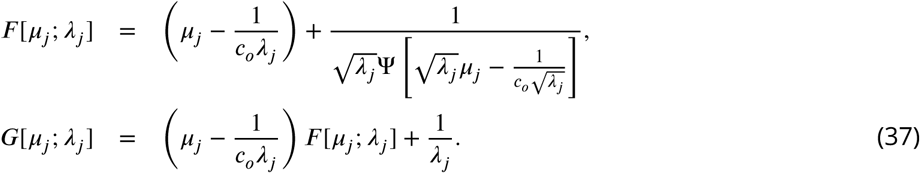

### 4.5 Learning of a concentration invariant representation

Up to now we focused on the expected concentration, ⟨*c*_*j*_⟩. However, in natural environments animals care more about whether or not an odor exists in its vicinity than what its concentration is. From a Bayesian perspective, this means the animals should compute the probability that an odor is present, denoted 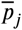. From Eq. (24), 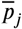 can be estimated as the steady state of the following dynamics,

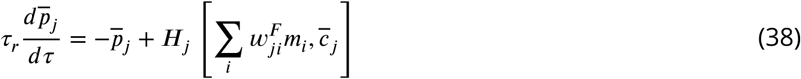

where ***H***_*j*_, which is approximately sigmoidal, is given, via Eq. (24),

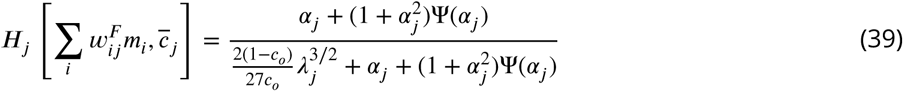

by with *α*_*j*_ given in Eq. (29).

In principle, neurons receiving input, *m*_*i*_, from M/T cells, such as layer 2 piriform cortex neurons, can decode the odor probability, as shown in Fig. 7A(i) and B(i). However, to calculate ***H***_*j*_ given input from M/T cells, the neuron would need to know the weights, 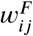, as well as *λ*_*j*_ and 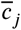 (the latter because *α*_*j*_ depends on 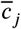; see Eq. (29)). This is clearly unrealistic, because there is no known biological mechanism that enables copying of the weights to the synapses on the neuron representing 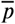. Moreover, because granule cells do not have output projections except for the dendro-dendritic connections with M/T cells, piriform neurons cannot know 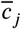 directly. Nevertheless, piriform neurons can learn to decode the concentration-invariant representation, 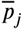, as follows.

Let us use 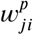 to denote the mean weight from M/T cells to the piriform neurons (see Fig. 7A(ii-iv)). Assume for the moment that 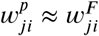; shortly we will write down a learning rule that achieves this. This takes care of the weight, but we also need an approximation to 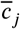. For that, we notice that if the estimation is unbiased, on average both 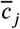 and 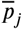 are equal to *c*_*o*_. Thus, the simplest way to approximate 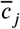 with the information available to the *j*^*th*^ piriform neuron is to use 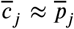. Under this approximation, and using 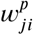 in place of 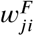, Eq. (38) becomes

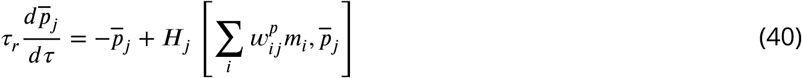

where ***H***_*j*_ is the same as Eq. (39), but with 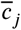 replaced by 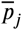. Finally, we make one more addition to the circuit, which is to add lateral inhibition, so 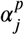 (the analog of *α*_*j*_, Eq. (29)) becomes

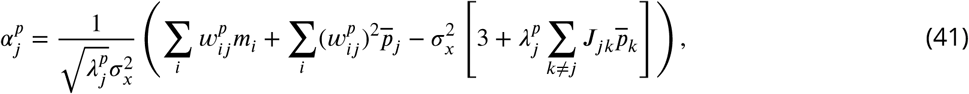

where, considering the analogy with Eq. (28b), 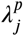 is given by

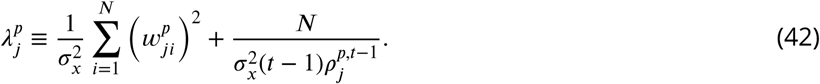

As above, 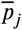 evolves with the weights set to their values updated at the end of previous trial. Once the neural dynamics reaches steady state, the weights are updated as in Eq. (34),

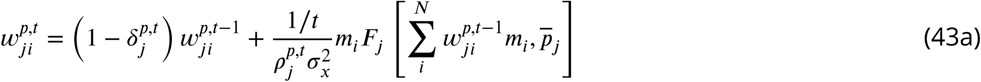

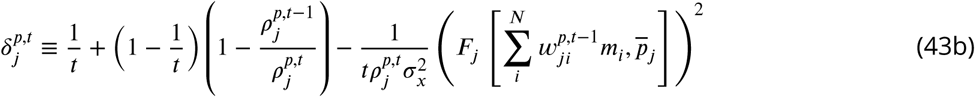

and the precision as in Eq. (27a),

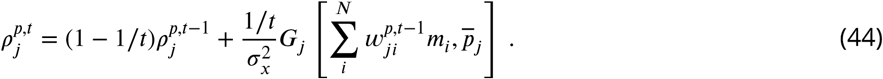

Here ***F***_*j*_ and ***G***_*j*_ are the estimated first/second moment given in Eq. (32) and Eq. (33), but calculated with 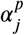 in Eq. (41). In addition, to ensure sparse piriform cell firing [Hiratani and Fukai, 2015], we introduced Hebbian plasticity to the lateral weights ***J***_*jk*_,

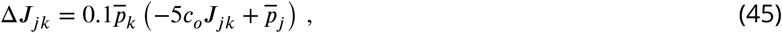

while bounding ***J***_*jk*_ > 0 and enforcing ***J***_*jj*_ = 0. We initialized ***J***_*jk*_ by ***J***_*jk*_ = 0.02.

In Fig. 7B(iv) and Fig. 7D-G, we modified the transfer function ***F***_*j*_ of granule cells by replacing the prior term *c*_*o*_ with the input from piriform neuron 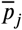. This means that 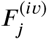 is written as

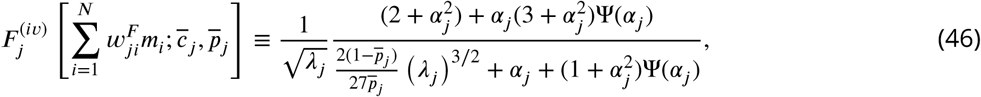

where *α*_*j*_ is still given by Eq. (29). We modulated *G*_*j*_ of granule cells, Eq. (33), in the same way, by replacing *c*_*o*_ with 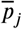. In Fig. 7C, we changed the relative strength of lateral inhibition simply by replacing ***J***_*jk*_ in Eq. (41) with [relative strength]***J***_*jk*_ where [relative strength] was set to 0 to 3, as shown in the x-axis of Fig. 7C, while using the original ***J***_*jk*_ for the weight update.

### 4.6 Reward-based Learning

Assuming that the reward amplitude depends only on the identity of odors, not on their concentrations, the reward, ***R***, on trial *t* is given by

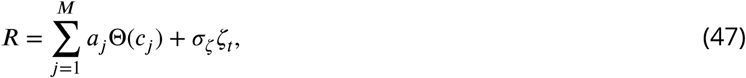

where *ζ*_*t*_ is a zero mean, unit variance Gaussian random variable.

To estimate the reward, we augment the circuit in Fig. 7A(iv) by introducing a set of neurons, denoted *e*_*p*_′ that receive input both from 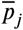 and the reward, ***R***; see Fig. 7E(i). Using *ā* to denote those weights, the natural neural dynamics of *e*_*p*_ is

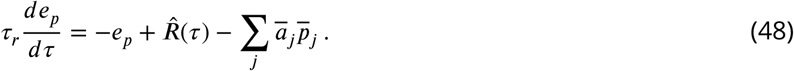

To represent the delay in reward delivery, 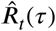 is zero for the first 2.5 seconds; after that it is set to the value of the reward,

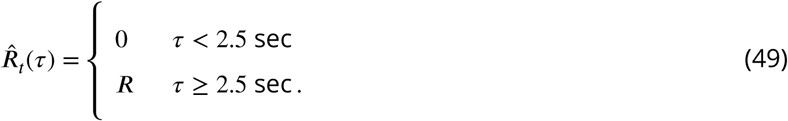

Note that for the first 2.5s of the trial, −*e*_*p*_ carries a prediction of the upcoming reward from the olfactory input, ***x***. Once the reward is provided, then the neuron represent the difference between the expected reward and the actual reward. That difference can be used to drive learning, via Hebbian plasticity,

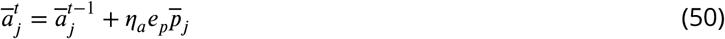

where *ā*_*j*_ is updated only after the reward has been presented. Importantly, *e*_*p*_ is evaluated after the reward presentation.

Similarly, for the direct readout from ***x*** depicted in Fig. 7E(ii), the reward is predicted by

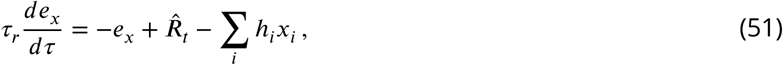

with *h*_*i*_ again update via Hebbian plasticity,

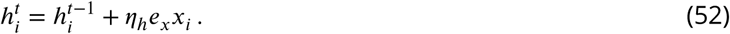

And again, *h*_*i*_ is not updated until the reward has been presented.

### 4.7 Sparse coding model

The sparse coding model originally proposed by Olshausen and colleagues [Olshausen and Field, 1996, 1997] can be applied to the model of olfactory learning as shown below. This model maximizes the likelihood of the data with respect to an unknown set of weights, denoted *ŵ*. The log-likelihood is given by

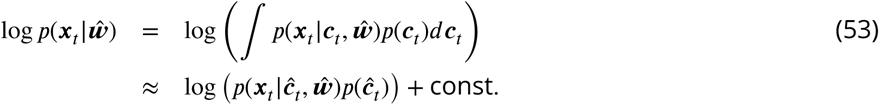

In the second line, the integral was approximated with the MAP estimate *ĉ*_*t*_ = argmax_***c***_ *p*(***x***_*t*_|***c***, *ŵ*)*p*(***c***). The objective function is thus given by

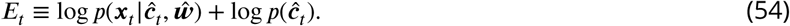

Because the noise on ***x***_*t*_ is Gaussian (see Eq. (6)), the first term is a simple quadratic function. However, the second term, log *p*(*ĉ*_*t*_), requires further approximation to remove the delta function, and thus ensure differentiability of *E*_*t*_ with respect to *ĉ*_*j*_. To this end, we approximated the prior with a Gamma distribution: 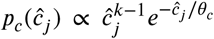. We used *k*_*c*_ = 3 and *0*_*c*_ = *c*_*o*_/3, ensuring a mean of *c*_*o*_. Under this approximation, the objective function, ***E***_*t*_, becomes

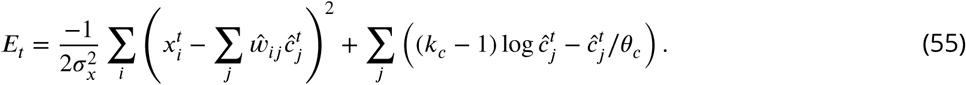

We maximize the objective function via stochastic gradient descent, which occurs in two steps. In the first step, we maximize *E*_*t*_ with respect to *ĉ*,

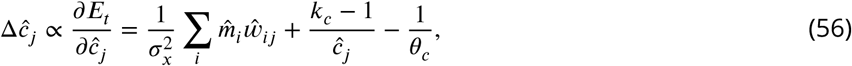

where 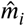 is the analog of Eq. (17),

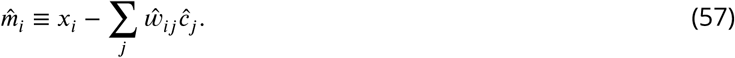

Once *ĉ*_*j*_ has converged, we update the weights via

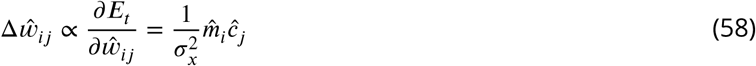

To prevent divergence of the weights, after each timestep we apply L-2 normalization (see below). In summary, on each trial, *t*, first, the *ĉ*_*j*_ (*J* = 1, 2, …, *M*) are updated,

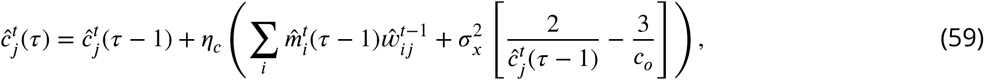

where the time step *τ* runs from 0 to 100000 in each trial. At the end of trial *t*, the weights are then updated by

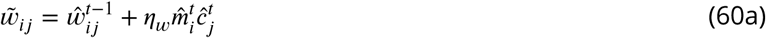

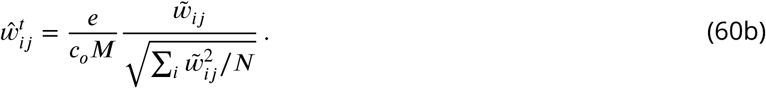

The learning rates, *η*_*c*_ and *η*_*w*_, were manually tuned. We used *η*_*c*_ = 0.00001 and *η*_*w*_ = 0.5 unless stated other-wise.

### 4.8 Simulation details

The parameters used in the simulations are given in Table 1. Additional details of the simulations are provided below. The main simulation codes are publicly available (http://github.com/nhiratani/olfactory_learning).

**Table 1:**
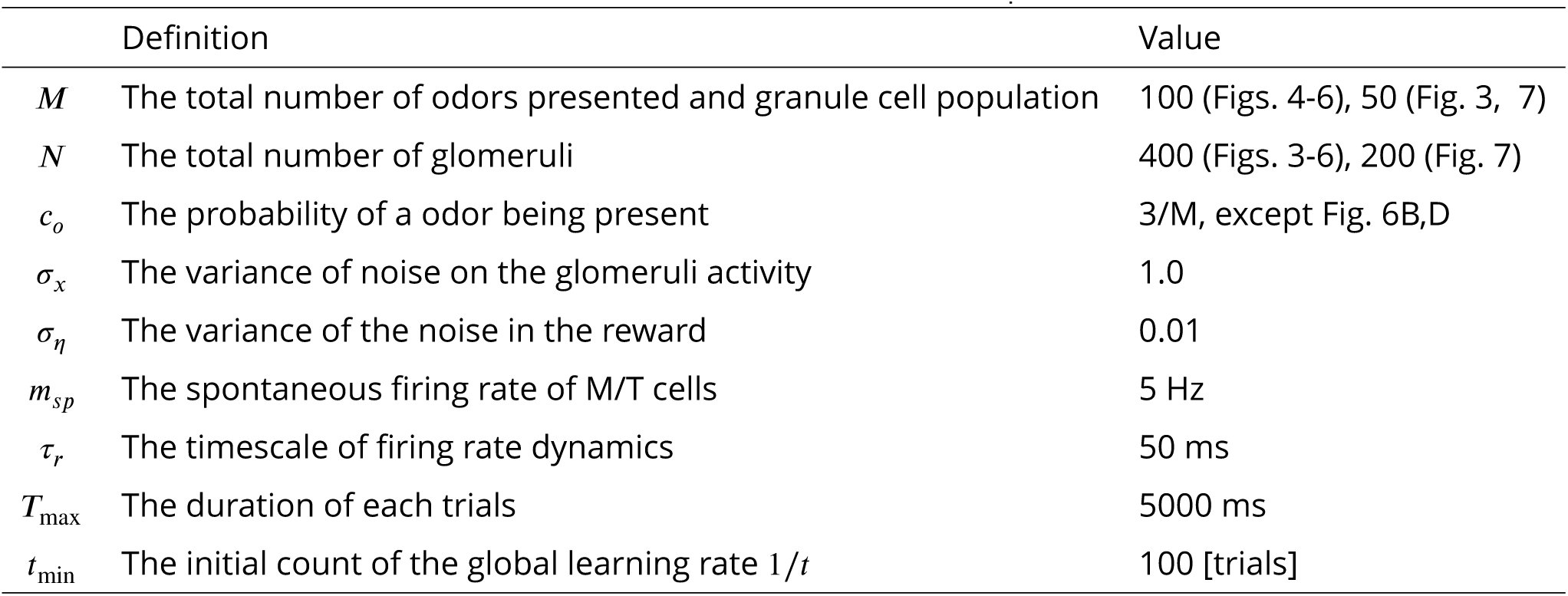
Definitions and values of the parameters

#### 4.8.1 Implementation of neural dynamics

The M/T cell activity, *m*_i_, was defined relative to a baseline, denoted *m*_*sp*_; in Fig. 3C, we plotted 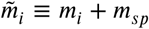. On each trial, *m*_*i*_ was initialized to zero and 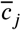 to 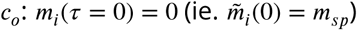 and 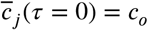. In addition, the firing rates were lower-bounded by *m*_*i*_ ≥ −*m*_*sp*_ and 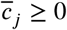.

To avoid numerical instability, Ψ(*α*) in Eq. (21b) was approximated as

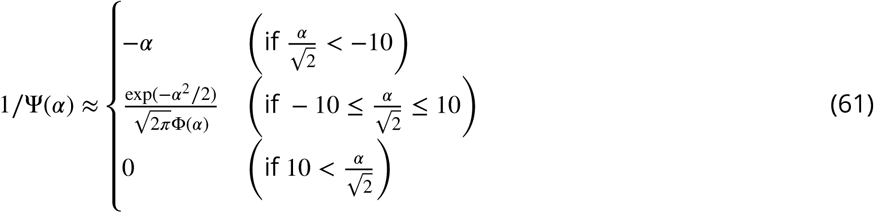

#### 4.8.2 Implementation of synaptic plasticity

Both the feedforward and the lateral weights were initially sampled from a lognormal distribution,

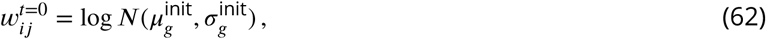

with the variance and mean parameters set to

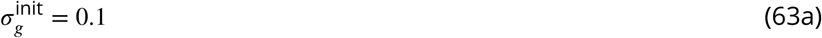

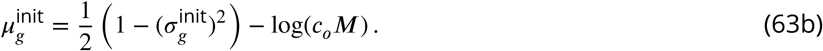

The precision factors, *ρ*_*j*_, were initialized as

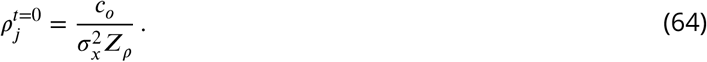

We used *Z*_*ρ*_ = 0.5, except in Figs. 6B and 6D, where we used *Z*_*ρ*_ = 0.3. The weights were lower-bounded by zero. As mentioned above, in the simulations we started *t* from *t* = *t*_min_ to suppress the influence of the initial samples.

#### 4.8.3 Learning with a fixed gain function

In Fig. 5D, we fixed all *λ*_*j*_ at 200.0 (gray) and 342.0 (black), while the 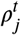 were updated at each trial as in Eq. (35).

#### 4.8.4 Learning with a fixed learning rate

Fixing the learning rate, 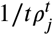 to a constant, denoted *η*, the learning rules for 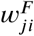 and 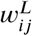 are rewritten as

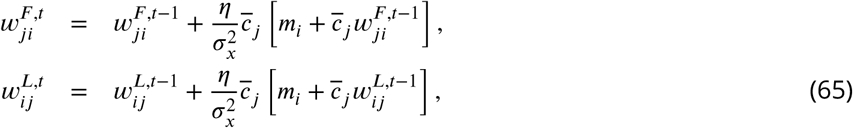

and *λ*_*j*_ is given by

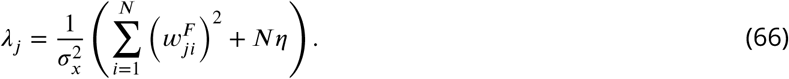

#### 4.8.5 Go/no go task

In the simulation of the go/no go task, we selected two odors (*j*_+_ and *j*_−_) out of ***M*** total odors, then randomly presented one or the other with concentrations drawn from a Gamma distribution (as in Eq. (8), but *c*_*j*_ > 0). The reward associated with *J*_+_ was ***R*** = 1.0 + *ζ* (i.e. 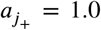), where *ζ* is the noise in the observed reward sampled from a zero-mean Gaussian with variance 0.01. The reward associated with *j*_−_ was ***R*** = *ζ* (i.e.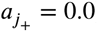).

Learning of the circuit shown in Fig.7E(i) was done in two steps. First, the weights, 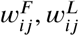 and 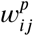, and the precisions, *ρ*_*j*_ and 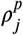, were learned with the unsupervised learning rules. During this unsupervised period, the reward, ***R***, was kept at zero. After 4000 trials of unsupervised learning, we fixed 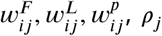, *and*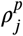, then trained the weights *ā*_*j*_ using Eq.50.

The reward weights for the circuits in both Figs. 7E(i) and 7E(ii), *ā*_*j*_ and *h*_*j*_, respectively, were initialized to zero, and the learning rates were manually tuned to the largest stable rates (*η*_*a*_ = 0.5 and *η*_*h*_ = 0.0015). The latter learning rate was smaller because ‖***x***‖ is typically much larger than 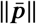, and also because the update of the *h*_*j*_ was more susceptible to instability.

The classification performance was measured by the probability that the predicted and actual reward were both above 0.5 or both below 0.5,

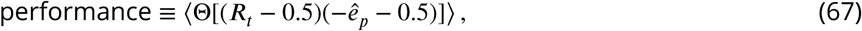

where *ê*_*p*_ is the value of *e*_*p*_ right before the reward delivery (*ê*_*p*_ = *e*_*p*_(*τ* = 2.45*s*)). Note that, as mentioned above, *ê*_*p*_ should converge to −***R***_*t*_, not ***R***_*t*_ in the given model. Thus, the average error was defined to be

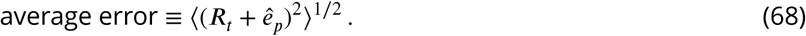

### 4.9 Performance evaluation

#### 4.9.1 Selectivity of granule cells

Because the network is trained with an unsupervised learning rule, we cannot know which neuron encodes which odor. We thus estimated the selectivity of a neuron from the incoming synaptic weights using a bootstrap method. Specifically, on each trial, the odor *o*(*j*) encoded by granule cell *j* is determined by choosing the odor that yields the maximum covariance between the estimated weights, ***w***^*F*^, and the true mixing weight, *w*,

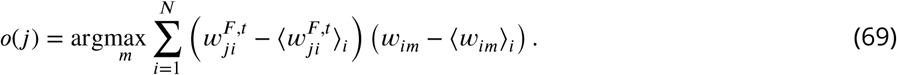

The selectivity can also be estimated from the activity of a neuron directly, and essentially the same result holds, although accurate readout of selectivity requires a large number of trials. After learning, most neurons learn to encode one odor stably.

#### 4.9.2 Odor estimation performance

Given the odor selectivity, *o*(*j*), the original odors can be reconstructed by

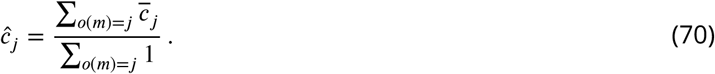

The denominator is the number of neurons that encode odor *j*, which converges to one after successful learning. If both the denominator and the numerator were zero, we set *ĉ*_*j*_ to 0. The performance was defined to be the correlation between the estimated odor concentration, *ĉ*_*j*_, and the true concentration, *c*_*j*_. Evaluation of performance on trial *t* used *o*(*j*) calculated from *w*^*F*, *t*−1^, not from *w*^*F*, *t*^. In Fig. 7B and C, we instead calculated the correlation between 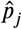 and the true value of Θ[*c*_*j*_] using the same method, where

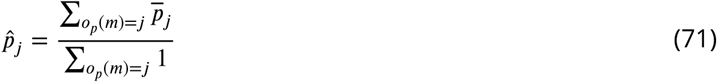

using the piriform neuron selectivity *o*_*p*_(*j*).

#### 4.9.3 Weight error

Given *o*(*j*), the error between the learned feedforward weight 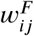 and the true mixing weight *w*_*ij*_ was calculated by

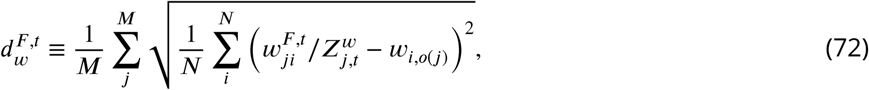

where 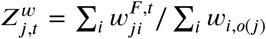. For comparison, in Fig. 6B, the weight errors were scaled by 7/3, so that the initial error was similar to the errors shown in Fig. 6A.

#### 4.9.4 Lifetime sparseness

For the measurement of the lifetime sparseness [Willmore and Tolhurst, 2001], we first presented individual odors *m* = 1, 2, …, ***M***, then recorded the activity of granule cells 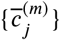. Subsequently, we calculated the sparseness by

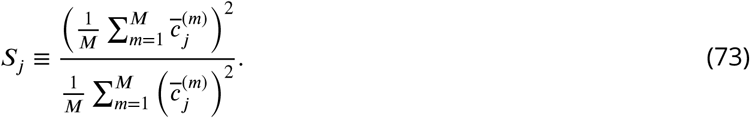

The lifetime sparseness, ***S***_*j*_, takes a small value (*S*_*j*_ ≃ 0) if the activity is sparse, while ***S***_*j*_ ≲ 1 is satisfied if the activity is uniform/homogeneous. Because of this, the lifetime sparseness is sometimes defined as 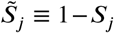 [Bolding and Franks, 2017].

## Acknowledgements

This work was supported by Gatsby Charitable Foundation and Wellcome Trust.

## References

Laurence Aitchison, Alex Pouget, and Peter E Latham. Probabilistic synapses. arXiv preprint 1410.1029, 2017.

Shunichi Amari. A theory of adaptive pattern classifiers. IEEE Transactions on Electronic Computers, 3:299–307, 1967.

Maxim Bazhenov, Ramon Huerta, and Brian H Smith. A computational framework for understanding decision making through integration of basic learning rules. Journal of Neuroscience, 33(13):5686–5697, 2013.

Matthew James Beal et al. Variational algorithms for approximate Bayesian inference. University of London London, 2003.

Jeff Beck, Alexandre Pouget, and Katherine A Heller. Complex inference in neural circuits with probabilistic population codes and topic models. In Advances in neural information processing systems, pages 3059–3067, 2012.

Kevin A Bolding and Kevin M Franks. Complementary codes for odor identity and intensity in olfactory cortex. Elife, 6:e22630, 2017.

Kevin A. Bolding and Kevin M. Franks. Recurrent cortical circuits implement concentration-invariant odor coding. Science, 361(6407), 2018. ISSN 0036-8075.

Alan Carleton, Leopoldo T Petreanu, Rusty Lansford, Arturo Alvarez-Buylla, and Pierre-Marie Lledo. Becoming a new neuron in the adult olfactory bulb. Nature neuroscience, 6(5):507, 2003.

Sophie JC Caron, Vanessa Ruta, LF Abbott, and Richard Axel. Random convergence of olfactory inputs in the drosophila mushroom body. Nature, 497(7447):113, 2013.

Frances S Chance, Larry F Abbott, and Alex D Reyes. Gain modulation from background synaptic input. Neuron, 35(4):773–782, 2002.

Xixi Chen, Li-Lian Yuan, Cuiping Zhao, Shari G Birnbaum, Andreas Frick, Wonil E Jung, Thomas L Schwarz, J David Sweatt, and Daniel Johnston. Deletion of kv4. 2 gene eliminates dendritic a-type k+ current and enhances induction of long-term potentiation in hippocampal ca1 pyramidal neurons. Journal of Neuroscience, 26(47):12143–12151, 2006.

Bradley B Doll, Dylan A Simon, and Nathaniel D Daw. The ubiquity of model-based reinforcement learning. Current opinion in neurobiology, 22(6):1075–1081, 2012.

Pierre Garrigues and Bruno A Olshausen. Learning horizontal connections in a sparse coding model of natural images. In Advances in Neural Information Processing Systems, pages 505–512, 2008.

Agnieszka Grabska-Barwińska, Simon Barthelmé, Jeff Beck, Zachary F Mainen, Alexandre Pouget, and Peter E Latham. A probabilistic approach to demixing odors. Nature neuroscience, 20(1):98, 2017.

Olivier Gschwend, Nixon M Abraham, Samuel Lagier, Frédéric Begnaud, Ivan Rodriguez, and Alan Carleton. Neuronal pattern separation in the olfactory bulb improves odor discrimination learning. Nature neuroscience, 18(10):1474, 2015.

Guillaume Hennequin, Yashar Ahmadian, Daniel B Rubin, Mate Lengyel, and Kenneth D Miller. The dynamical regime of sensory cortex: stable dynamics around a single stimulus-tuned attractor account for patterns of noise variability. Neuron, 98(4):846–860, 2018.

Naoki Hiratani and Tomoki Fukai. Mixed signal learning by spike correlation propagation in feedback in-hibitory circuits. PLoS computational biology, 11(4):e1004227, 2015.

Naoki Hiratani and Tomoki Fukai. Redundancy in synaptic connections enables neurons to learn optimally. Proceedings of the National Academy of Sciences, 115(29):E6871–E6879, 2018.

JJ Hopfield. Odor space and olfactory processing: collective algorithms and neural implementation. Proceedings of the National Academy of Sciences, 96(22):12506–12511, 1999.

Aapo Hyvärinen and Erkki Oja. Independent component analysis: algorithms and applications. Neural net-works, 13(4-5):411–430, 2000.

Kentaro K Ishii, Takuya Osakada, Hiromi Mori, Nobuhiko Miyasaka, Yoshihiro Yoshihara, Kazunari Miyamichi, and Kazushige Touhara. A labeled-line neural circuit for pheromone-mediated sexual behaviors in mice. Neuron, 95(1):123–137, 2017.

Daniel Kepple, Brittany N Cazakoff, Heike S Demmer, Dennis Eckmeier, Stephen D Shea, and Alexei A Koulakov. Computational algorithms and neural circuitry for compressed sensing in the mammalian main olfactory bulb. bioRxiv, page 339689, 2018.

Alexei A Koulakov and Dmitry Rinberg. Sparse incomplete representations: a potential role of olfactory granule cells. Neuron, 72(1):124–136, 2011.

Qian Li and Stephen D Liberles. Aversion and attraction through olfaction. Current Biology, 25(3):R120–R129, 2015.

Christiane Linster, Brett A Johnson, Alix Morse, Esther Yue, and Michael Leon. Spontaneous versus reinforced olfactory discriminations. Journal of Neuroscience, 22(16):6842–6845, 2002.

David JC MacKay. A practical bayesian framework for backpropagation networks. Neural computation, 4(3): 448–472, 1992.

Alexander Mathis, Dan Rokni, Vikrant Kapoor, Matthias Bethge, and Venkatesh N Murthy. Reading out olfactory receptors: feedforward circuits detect odors in mixtures without demixing. Neuron, 91(5):1110–1123, 2016.

Guo-li Ming and Hongjun Song. Adult neurogenesis in the mammalian brain: signi1cant answers and significant questions. Neuron, 70(4):687–702, 2011.

Toby J Mitchell and John J Beauchamp. Bayesian variable selection in linear regression. Journal of the American Statistical Association, 83(404):1023–1032, 1988.

Vinod Nair and Geoffrey E Hinton. Rectified linear units improve restricted boltzmann machines. In Proceedings of the 27th international conference on machine learning (ICML-10), pages 807–814, 2010.

Antoine Nissant, Cedric Bardy, Hiroyuki Katagiri, Kerren Murray, and Pierre-Marie Lledo. Adult neurogenesis promotes synaptic plasticity in the olfactory bulb. Nature neuroscience, 12(6):728, 2009.

Robert J O’Connell and Maxwell M Mozell. Quantitative stimulation of frog olfactory receptors. Journal of neurophysiology, 32(1):51–63, 1969.

Bruno A Olshausen and David J Field. Emergence of simple-cell receptive field properties by learning a sparse code for natural images. Nature, 381(6583):607, 1996.

Bruno A Olshausen and David J Field. Sparse coding with an overcomplete basis set: A strategy employed by v1? Vision research, 37(23):3311–3325, 1997.

Anne-Marie M Oswald and Alex D Reyes. Maturation of intrinsic and synaptic properties of layer 2/3 pyramidal neurons in mouse auditory cortex. Journal of neurophysiology, 99(6):2998–3008, 2008.

Panayiota Poirazi, Terrence Brannon, and Bartlett W Mel. Pyramidal neuron as two-layer neural network. Neuron, 37(6):989–999, 2003.

Benjamin Roland, Thomas Deneux, Kevin M Franks, Brice Bathellier, and Alexander Fleischmann. Odor identity coding by distributed ensembles of neurons in the mouse olfactory cortex. Elife, 6:e26337, 2017.

Mala M Shah, Rebecca S Hammond, and Dax A Hoffman. Dendritic ion channel traZcking and plasticity. Trends in neurosciences, 33(7):307–316, 2010.

Gordon M Shepherd, Wei R Chen, David Willhite, Michele Migliore, and Charles A Greer. The olfactory granule cell: from classical enigma to central role in olfactory processing. Brain research reviews, 55(2):373–382, 2007.

U Staubli, D Fraser, R Faraday, and G Lynch. Olfaction and the” data” memory system in rats. Behavioral neuroscience, 101(6):757, 1987.

Jie Tan, Agnès Savigner, Minghong Ma, and Minmin Luo. Odor information processing by the olfactory bulb analyzed in gene-targeted mice. Neuron, 65(6):912–926, 2010.

Sina Tootoonian and Máté Lengyel. A dual algorithm for olfactory computation in the locust brain. In Advances in Neural Information Processing Systems, pages 2276–2284, 2014.

Jochen Triesch. Synergies between intrinsic and synaptic plasticity in individual model neurons. In Advances in neural information processing systems, pages 1417–1424, 2005.

Jenelle L Wallace, Martin Wienisch, and Venkatesh N Murthy. Development and re1nement of functional properties of adult-born neurons. Neuron, 96(4):883–896, 2017.

Daniel W Wesson and Donald A Wilson. SniZng out the contributions of the olfactory tubercle to the sense of smell: hedonics, sensory integration, and more? Neuroscience & Biobehavioral Reviews, 35(3):655–668, 2011.

Benjamin Willmore and David J Tolhurst. Characterizing the sparseness of neural codes. Network: Computation in Neural Systems, 12(3):255–270, 2001.

Rachel I Wilson and Zachary F Mainen. Early events in olfactory processing. Annu. Rev. Neurosci., 29:163–201, 2006.

Yoshiyuki Yamada, Khaleel Bhaukaurally, Tamás J Madarász, Alexandre Pouget, Ivan Rodriguez, and Alan Carleton. Context-and output layer-dependent long-term ensemble plasticity in a sensory circuit. Neuron, 93(5):1198–1212, 2017.

Zhong-wei Zhang. Maturation of layer v pyramidal neurons in the rat prefrontal cortex: intrinsic properties and synaptic function. Journal of neurophysiology, 91(3):1171–1182, 2004.

